# Optimization of the fluorogen-activating protein tag for quantitative protein trafficking and co-localization studies in *S. cerevisiae*

**DOI:** 10.1101/2024.04.20.590399

**Authors:** Katherine G. Oppenheimer, Natalie A. Hager, Ceara K. McAtee, Elif Filiztekin, Chaowei Shang, Justina A. Warnick, Marcel P. Bruchez, Jeffrey L. Brodsky, Derek C. Prosser, Adam V. Kwiatkowski, Allyson F. O’Donnell

## Abstract

Spatial and temporal tracking of fluorescent proteins in live cells permits visualization of proteome remodeling in response to extracellular cues. Historically, protein dynamics during trafficking have been visualized using constitutively active fluorescent proteins (FPs) fused to proteins of interest. While powerful, such FPs label all cellular pools of a protein, potentially masking the dynamics of select subpopulations. To help study protein subpopulations, bioconjugate tags, including the fluorogen activation proteins (FAPs), were developed. FAPs are comprised of two components: a single-chain antibody (SCA) fused to the protein of interest and a malachite-green (MG) derivative, which fluoresces only when bound to the SCA. Importantly, the MG derivatives can be either cell-permeant or -impermeant, thus permitting isolated detection of SCA-tagged proteins at the cell surface and facilitating quantitative endocytic measures. To expand FAP use in yeast, we optimized the SCA for yeast expression, created FAP-tagging plasmids, and generated FAP-tagged organelle markers. To demonstrate FAP efficacy, we coupled the SCA to the yeast G-protein coupled receptor Ste3. We measured Ste3 endocytic dynamics in response to pheromone and characterized *cis*- and *trans*-acting regulators of Ste3. Our work significantly expands FAP technology for varied applications in *S. cerevisiae*.

**SIGNIFICANCE STATEMENT:**

- Quantitative endocytic assays are required to characterize factors that regulate both ligand-dependent and constitutive endocytosis.
- We optimize fluorogen-activating proteins (FAPs) technology for use as a live cell imaging probe in yeast that fluoresces in the far-red range for quantitative endocytosis assays.
- The FAP tools and approaches generated will facilitate quantitative endocytic and protein recycling assays for yeast cell biologists.

## INTRODUCTION

Eukaryotic cells control the distribution of proteins within membrane-bound organelles via selective protein trafficking. At the plasma membrane (PM), protein abundance is regulated by both exocytic and endocytic events. Achieving the correct balance of PM proteins is critical for nutrient homeostasis and dynamic cellular responses, such as adaptation to stress, hormones, or other signaling molecules (Sorkin and Von Zastrow, 2002; O’Donnell and Schmidt, 2019; Hager *et al*., 2021). Defective membrane trafficking or aberrant activity of membrane proteins at the PM cause disease, including diabetes, cardiac arrhythmias, and Alzheimer’s (Aridor and Hannan, 2000; Howell *et al*., 2006; Hung and Link, 2011; Yarwood *et al*., 2020). To better define how defective protein trafficking contributes to these and other disorders, it is necessary to first distinguish changes in protein activity from alterations in protein localization (Perkins and Bruchez, 2020). For example, mutations affecting the synthesis, folding, trafficking, or activity (i.e., gating or conductance) of the cystic fibrosis transmembrane conductance regulator (CFTR) all cause cystic fibrosis. Still, unique therapies that improve CFTR function are only effectively deployed when the underlying molecular mechanism for disease is understood (Moskowitz *et al*., 2008; Koulov *et al*., 2010; Coppinger *et al*., 2012; Lopes-Pacheco, 2016, 2019). Quantitative studies of endocytosis and intracellular protein sorting are imperative to identify and characterize mutations that compromise protein trafficking.

Techniques such as cell-surface biotinylation assays and ligand labeling are frequently used to quantify protein trafficking to and from the PM, yet both have their limitations (Nishimura and Sasaki, 2008; Bohme and Beck-Sickinger, 2009; Chen *et al*., 2011; Tham and Moukhles, 2017). For example, cell-surface biotinylation can abnormally trigger endocytosis (Dundas *et al*., 2013) and may underestimate the abundance of single-pass membrane proteins (Tham and Moukhles, 2017). In addition, biotinylation of surface proteins is ineffective in some cell types and model systems. For example, in yeasts like *Saccharomyces cerevisiae* and *Candida albicans,* avidin non-specifically binds the cell wall (Masuoka *et al*., 2002), while in select human cell types, some membrane proteins associate non-specifically with streptavidin (Cole *et al*., 1987). Furthermore, extracting biotinylated membrane proteins can be problematic; in yeast and plants, the cell wall provides an added barrier to effective membrane protein solubilization and isolation (Smith *et al*., 2000; Klis *et al*., 2002; Powell *et al*., 2003; Mukherjee *et al*., 2020). Likewise, quantitative endocytic assays using receptor-specific ligand probes also have limitations and require selective derivatization of each ligand.

The modified ligands must also be PM-impermeant, bind the receptor similar to native ligand, and remain bound to monitor internalization and post-endocytic sorting (Los *et al*., 2008; Komatsu *et al*., 2011; Leng *et al*., 2017; Jonker *et al*., 2020). Radiolabeled or rhodamine-labeled epidermal growth factor (EGF) has been a powerful tool for studying the endocytosis, sorting, and recycling of the epidermal growth factor receptor (EGFR) in many cell types and models (Huang *et al*., 2007; Goh *et al*., 2010; Tomas *et al*., 2014; Tanaka *et al*., 2018). Similarly, in *S. cerevisiae*, radiolabeled α-factor, the mating pheromone that binds to the G-protein-coupled receptor (GPCR) Ste2, was initially used to study Ste2 endocytic dynamics (Zanolari and Riezman, 1991; Hicke *et al*., 1998; Dunn and Hicke, 2001; Shih *et al*., 2002; Toshima *et al*., 2009). Later, fluorescently-tagged α-factor was used to stimulate Ste2 internalization and illuminate its ligand-induced trafficking (Toshima *et al*., 2006). Although useful, these tools are not easily adapted to explore the many proteins trafficked to and from the PM.

In addition to cell-surface biotinylation and ligand labeling, fluorescent proteins (FPs) such as the green fluorescent protein (GFP) and its derivatives are widely used to monitor membrane protein dynamics (Lippincott-Schwartz *et al*., 2003; Miyawaki *et al*., 2003). Despite the wealth of applications for FPs and the bounty of available probes, FPs have some shortcomings. For example, high fluorescence levels in intracellular compartments can sometimes prevent clear differentiation of the PM pool of a tagged protein. This intracellular fluorescence can complicate quantitative measures of endocytic turnover or PM delivery. While the use of pH-sensitive FP derivatives, such as the GFP derivative pHluorin, facilitates quantitative endocytic assays by quenching the fluorescence of intracellular sub-populations (Balaji and Ryan, 2007; Prosser *et al*., 2010; Nicholson-Fish *et al*., 2016; Prosser *et al*., 2016), this approach requires transit to an acidic organelle, e.g., the multi-vesicular body (MVB) or lysosome (yeast vacuole equivalent). Furthermore, these pH-sensitive probes are ineffective in recycling assays that assess the return of endocytosed membrane proteins to the PM.

As an alternative to FPs, bioconjugate tags can permit temporal and spatial fluorescent labeling and simultaneous visualization of specific proteins. Unlike FPs, some bioconjugate tags form covalent bonds with chemical fluorophores (Griffin *et al*., 1998); thus, light is emitted when a peptide tag binds to the probe. However, these probes exhibit background fluorescence due to their affinity for cysteine-rich proteins (Griffin *et al*., 1998; Zurn *et al*., 2010; Gallo, 2020). In contrast, appended Halo, CLIP, SNAP, LAP, and BL motifs engineered onto proteins of interest form irreversible bonds with fluorescently labeled ligands (Keppler *et al*., 2003; Gautier *et al*., 2008; Los *et al*., 2008; Watanabe *et al*., 2010; Yao *et al*., 2012). These tags then fluoresce when bound to an added dye, which has advantages for spatial and temporal resolution (Griffin *et al*., 1998; Keppler *et al*., 2003; Keppler *et al*., 2004; Chen *et al*., 2005; Fernandez-Suarez *et al*., 2007; Luedtke *et al*., 2007; Gautier *et al*., 2008; Gallo, 2020).

Another system for bioconjugate tagging is the use of fluorogen-activating protein (FAP) technology, which offers selective labeling of specific membrane protein pools, enhanced spatiotemporal visualization, and added flexibility in imaging parameters and approaches (Szent-Gyorgyi *et al*., 2008; Boeck and Spencer, 2017; Emmerstorfer-Augustin *et al*., 2018; Hager *et al*., 2018; Gallo, 2020; Perkins and Bruchez, 2020). FAP tags are comprised of a genetically encoded single-chain antibody (SCA) fused to a protein of interest. The variable region of the SCA displays a high affinity for synthetic compounds, *i.e.*, fluorogens. When free, neither the SCA nor the fluorogen are fluorescent (Figure 1) (Szent-Gyorgyi *et al*., 2008; Fisher *et al*., 2010; Gallo, 2020; Perkins and Bruchez, 2020). However, when the fluorogen binds the SCA, the fluorogen conformation is restricted, resulting in a significant (up to 20,000-fold) increase in fluorescence (Lee *et al*., 1986; Silva *et al*., 2007; Shank *et al*., 2013; Gallo, 2020), matching the sensitivity and intensity of conventional FPs such as EGFP and mCherry (Szent-Gyorgyi *et al*., 2013). Based on this exceptional increase in fluorescence, FAPs also offer a better signal-to-noise ratio than FPs, and the dye does not need to be washed away for certain applications (Perkins and Bruchez, 2020). FAP emission wavelengths are also wide-ranging and adaptable to a variety of filter sets, since the fluorogens can be derived from several dyes, including malachite green (MG). Another key feature of FAP technology is the ability to use either cell-permeant or -impermeant fluorogens (Figure 1, B and C). In mammalian cells, FAPs have been used to examine endocytosis, sorting, and recycling of GABA_A_ receptors in neurons (Lorenz-Guertin *et al*., 2017), endocytosis of β_2_-adrenergic receptors (β_2_AR) in NIH3T3 fibroblasts (Holleran *et al*., 2010), β_2_AR shuttling to and from the PM in lymphoma cells (Fisher *et al*., 2010), and mutant CFTR trafficking to the PM in human bronchial epithelial cells and HEK293T cells (Holleran *et al*., 2012; Goeckeler-Fried *et al*., 2021). Further, permeant fluorogens can be used in conjunction with impermeant fluorogens to reveal total cellular protein versus the cell surface pool, shedding light on changes in protein turnover and trafficking to a specific organelle (Pratt *et al*., 2015; Yan *et al*., 2015; Naganbabu *et al*., 2016; Perkins and Bruchez, 2020). FAP technology can also be adapted to measure PM recycling rates when two differentially fluorescent fluorogens are used (Szent-Gyorgyi *et al*., 2013; Shiwarski *et al*., 2017).

**Figure 1:**
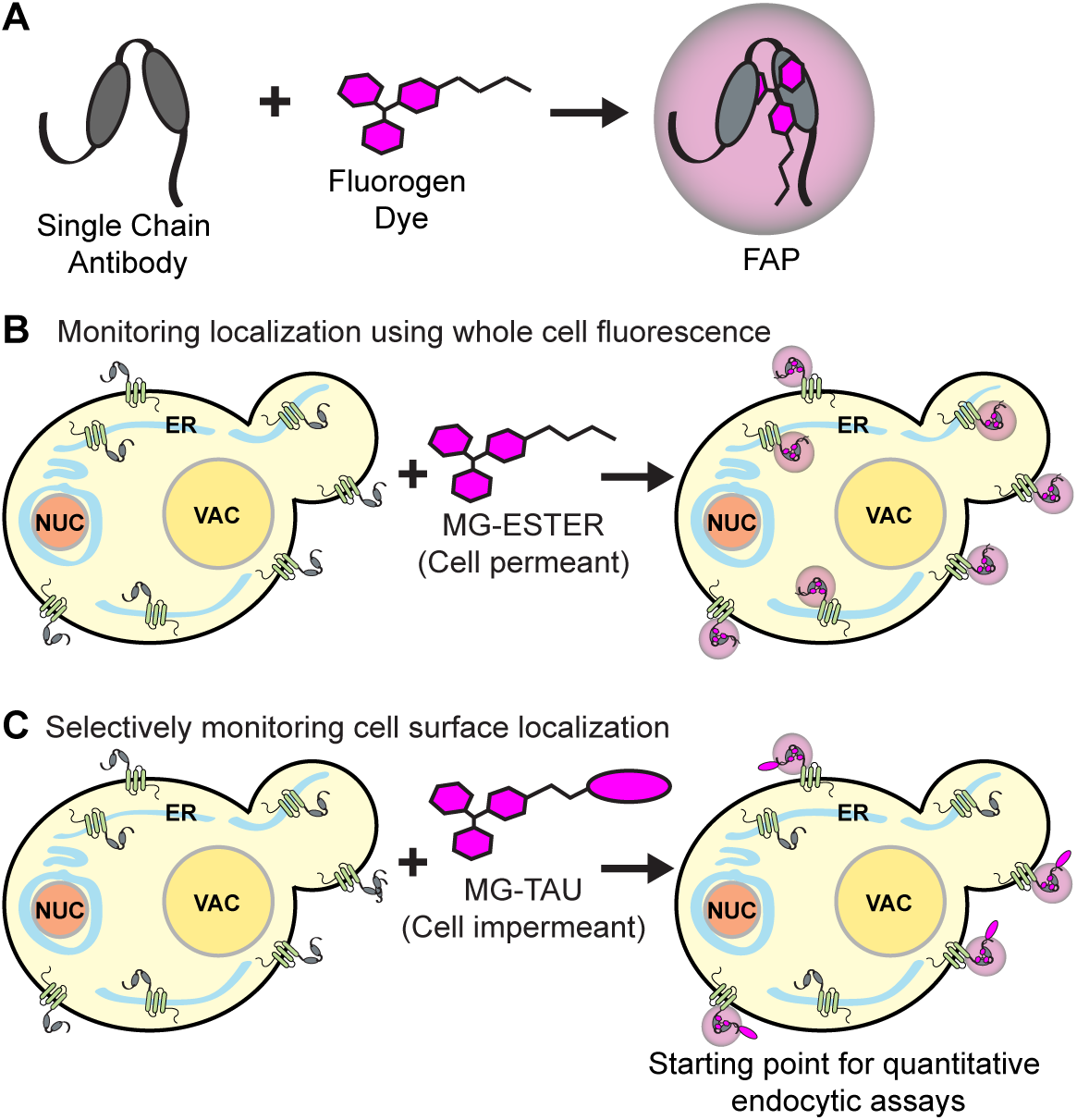
Model of FAP technology. (A) A single-chain antibody (SCA) binds to the fluorogen dye, locking it into a conformation that promotes fluorescence. (B-C) The fluorogen dye can be cell-permeant (*i.e*., MG-ESTER) or -impermeant (*i.e*., MG-TAU). While MG-ESTER allows for visualization of the entire pool of SCA-tagged protein, MG-TAU allows for selective visualization of the cell surface pool. By adding dye for a brief time and washing excess away, one can quantitively monitor endocytosis of the PM-labeled pool from the cell surface.

Despite its extensive uses, the FAP technology has not been widely adapted for use in the budding yeast *Saccharomyces cerevisiae* even though SCAs were developed as a result of yeast screens (Feldhaus *et al*., 2003; Szent-Gyorgyi *et al*., 2008). Nevertheless, we and others have recently used FAP technology to track the sorting of membrane proteins successfully (Emmerstorfer-Augustin *et al*., 2018; Hager *et al*., 2018). Here, we significantly expand on these initial studies by enhancing the performance and accessibility of FAP technology in yeast. We codon-optimized the SCA for expression in *S. cerevisiae* and confirmed that optimized FAP exhibits increased stability. We then created an extensive collection of FAP-tagged cloning vectors for yeast and built a suite of FAP-tagged cellular markers for co-localization studies in yeast. We then demonstrated the utility of the FAP-tagging approach by characterizing *cis*- and *trans*-acting regulators of the ligand-induced endocytosis of Ste3, the GPCR that controls mating pathway signaling in Mat α yeast cells (Bardwell *et al*., 1994; Bardwell, 2005). We expect these new FAP tools to be a rich, widely applicable resource for the yeast cell biology community. FAP-tagged Ste3 and other membrane cargos will allow us to define the factors needed for post-endocytic recycling in future studies.

## RESULTS

### FAP codon optimization for yeast expression

We first codon optimized the SCA motif (described as dL5 in (Szent-Gyorgyi *et al*., 2008; Yan *et al*., 2015) for expression in *Saccharomyces cerevisiae* using the JAVA Codon Adaptation Tool (JCat) (Grote *et al*., 2005). In JCat, the Codon Adaption Index (CAI) ranges from 0 to 1, where values approaching 1 indicate a codon with an abundant tRNA in that organism. Before optimization, original FAP (FAP_origin_) had an average CAI of 0.063, indicating that most codons were matched to low abundance yeast tRNAs. In contrast, the codon-optimized FAP (FAP_optim_) sequence had a CAI value of 0.973 (Figure 2A). We also introduced an N- or C-terminal linker (sequence Ala-Gly-Ala-Gly-Ala-Gly) to facilitate FAP folding independently of the protein to which it was fused, and a MYC epitope to facilitate biochemical detection (Figure 2B).

**Figure 2:**
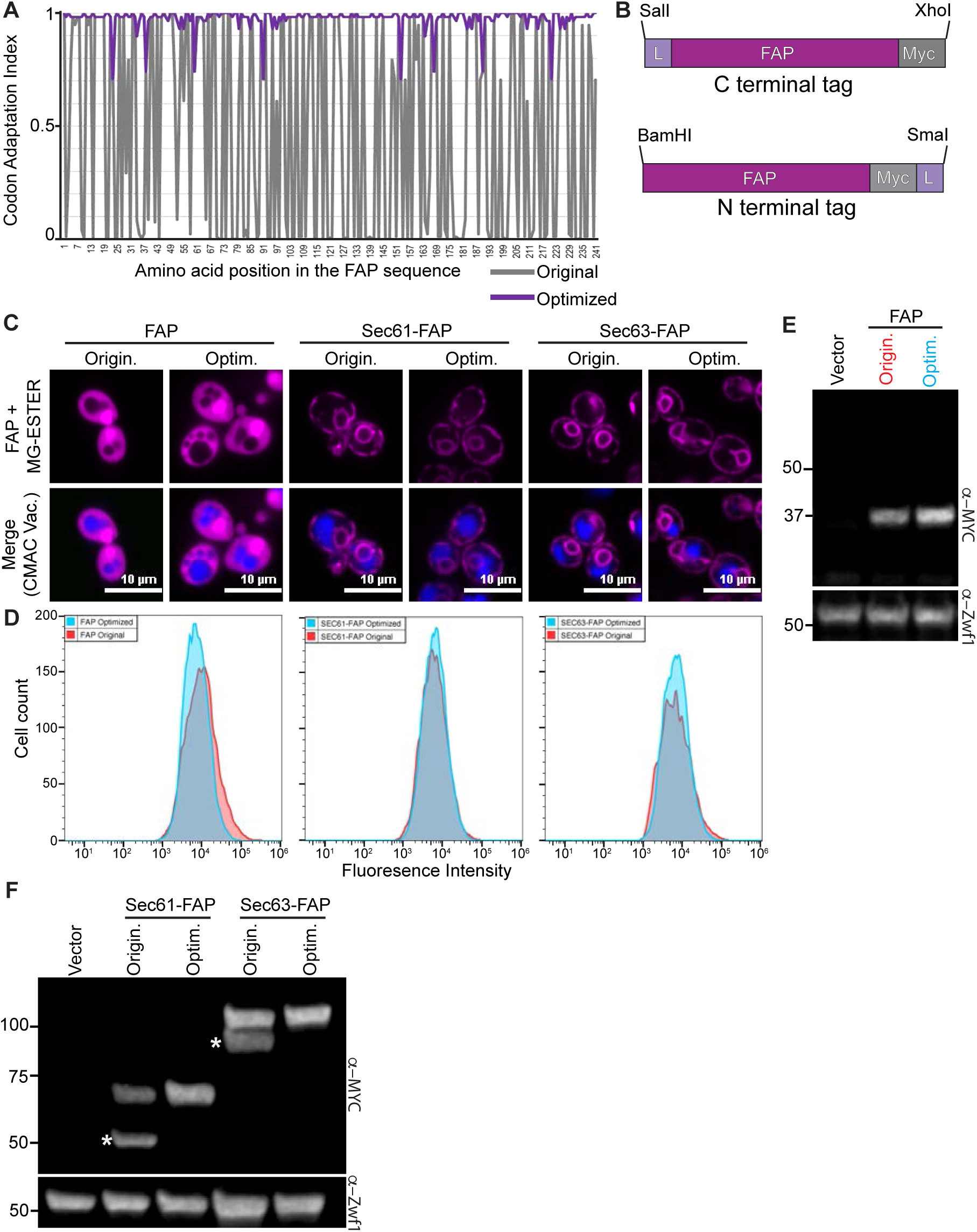
Codon optimization of the FAP sequence for expression in yeast. (A) Graph of the codon adaptation index for *S. cerevisiae* of the Original FAP sequence (grey) versus the codon-optimized FAP sequence (purple). (B) Schematic of the N- or C-terminal FAP tagging cassettes. L = linker (Ala-Gly)_3_ and Myc = 2xMYC epitope tag. (C) Confocal fluorescence microscopy of yeast expressing soluble FAP, Sec61- or Sec63-FAP, either as the original (Origin.) or the codon-optimized FAP sequences (Optim.; magenta). Vacuoles are stained with CMAC (blue) in the merge. (D) Histograms of flow cytometry data showing fluorescence distributions for cells expressing the different FAP-fusions from panel C. (E-F) Immunoblots of yeast extracts from cells expressing an empty vector control, and either (E) soluble FAP or (F) FAP-tagged Sec61 or Sec63 as original or optimized versions. Zwf1 is the loading control. The white asterisks in (F) identify breakdown products observed for the FAP_Origin_ tagged proteins. Numbers on the left represent MW markers in kDa.

We then sought to determine whether the FAP_optim_ improved expression and function in yeast. Both FAP_optim_ and FAP_origin_ were expressed under the control of the *TEF1* promoter as a soluble protein (“FAP”) or as fusions to the ER-resident membrane proteins Sec61 and Sec63. Using flow cytometry and fluorescence microscopy, we found no difference in the fluorescence intensity of FAP_optim_ compared to FAP_origin_ regardless of whether they were expressed as free soluble or fusion proteins (Figure 2, C and D and Supplemental Figure 1, A and B). While free FAP_origin_ and FAP_optim_ were roughly equivalent on immunoblots, when the FAP-tagged fusion proteins were examined via immunoblotting, we consistently observed breakdown products with FAP_origin_-fused proteins (Figure 2, E-F). For Sec61- and Sec63-FAP_origin_, we observe bands at the expected molecular masses for intact fusions (75 kDa and 105 kDa, respectively), but significant breakdown products were also detected (∼50 kDa and 90 kDa, respectively) (Figure 2F). Since the MYC tag detected in these Sec61- and Sec63-FAP_origin_ fusions is at the C-terminus of the protein, these breakdown products must be due to cleavage within the Sec61 or Sec63 proteins themselves and not in the FAP tag, which is only 24.2 kDa. Importantly, these breakdown products were absent with Sec61- and Sec63-FAP_optim_ fusions (Figure 2F), suggesting that proteins tagged with FAP_origin_ are more susceptible to cleavage and degradation. If present in live cells, these degradation products could lead to spurious determinations of protein localization. Therefore, all subsequent studies in our work used the FAP_optim_ moiety.

### Cellular growth conditions and vacuolar proteases influence FAP fluorescence

Next, we explored conditions that might impact FAP_optim_ signal intensity. The FAP_origin_ signal is reportedly influenced by changes in pH, with reduced fluorescence at the cell surface of the Ste2 receptor in acidic environments and an optimal fluorescent signal at pH 6.5 (Emmerstorfer-Augustin *et al*., 2018). We found that soluble, intracellular FAP_optim_ and Sec61-FAP_optim_ were unaffected by pH changes, with equivalent fluorescence intensities observed in cells incubated at pHs ranging from 4.1 to 7.0 by both confocal microscopy and flow cytometry (Figure 3, A-C and Supplemental Figure 2, A-E). These data indicate that intracellular FAP_optim_ probes can be used in yeast cells grown in media at a wide range of pHs. This may be partly due to the fact that even yeast cells grown in acidic conditions (∼pH 4.0) can buffer their cytosols to maintain a near-optimal intracellular pH of 6 (Valli *et al*., 2005).

**Figure 3:**
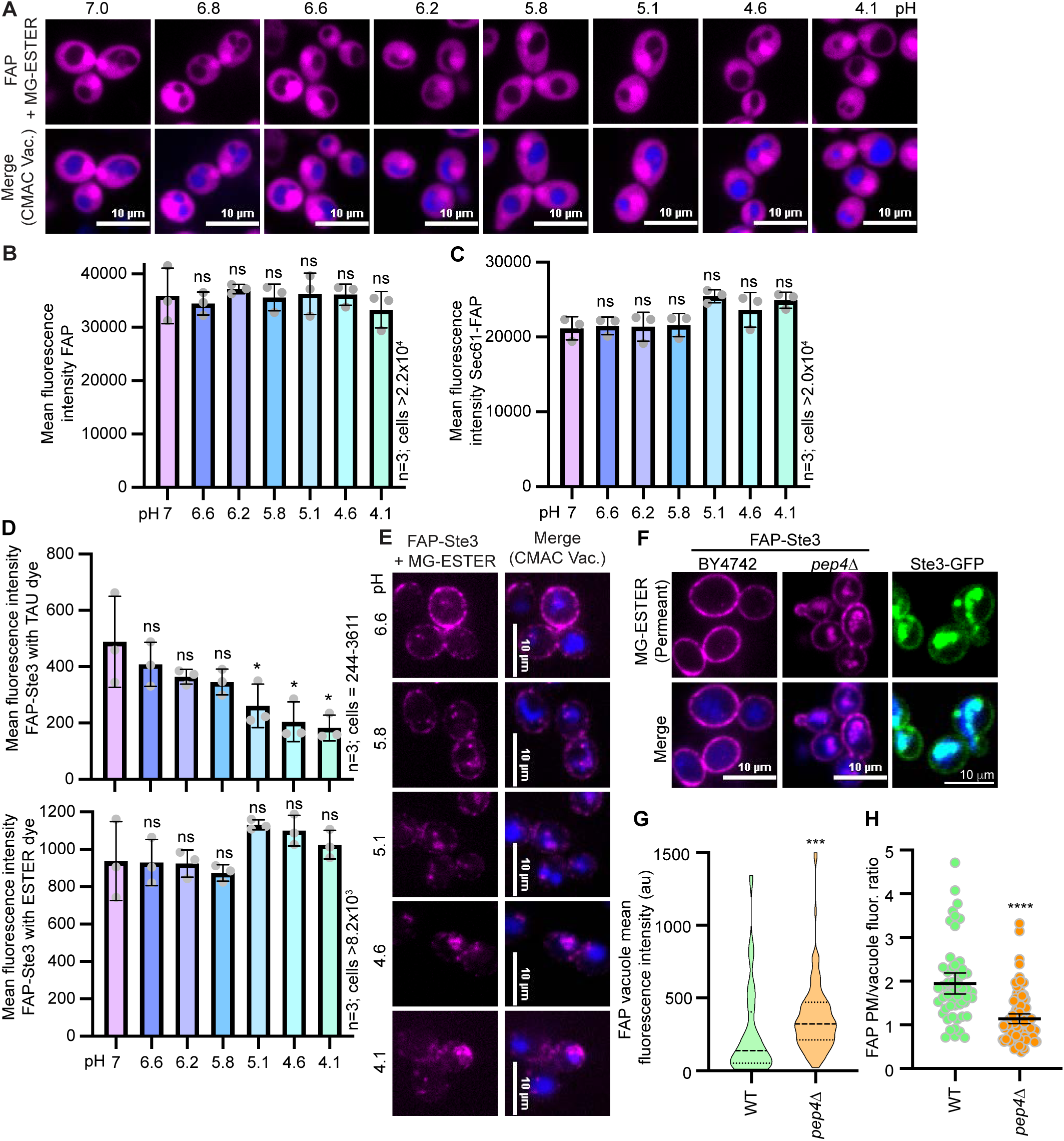
Parameters that impact FAP fluorescence in yeast. (A and E) Confocal fluorescence microscopy images of cells expressing (A) soluble FAP_OPTIM_ or (E) FAP_OPTIM_-Ste3. Cells were incubated in different pH media for 2 h before adding MG-ESTER (magenta) and CMAC (blue) dyes and imaging. (B-C) Bar graphs representing the mean of three biological replicates of flow cytometry data for cells as shown in panel A (in panel B) or those expressing FAP-Sec61 (in panel C). ANOVA statistical test assessed changes in fluorescence between samples relative to pH 7.0 (ns = not significant). (D) Bar graphs of the mean FAP-Ste3 fluorescence intensities with MG-TAU (top) or -ESTER (bottom) dyes from three replicate flow cytometry experiments are shown. ANOVA statistical test assessed changes in fluorescence between samples relative to pH 7.0 (ns = not significant; * p<0.05). (F) Confocal fluorescence microscopy of cells expressing either FAP-Ste3 (magenta) or Ste3-GFP (green) in the indicated genetic backgrounds. (G-H) Quantification of the vacuole signal intensity (violin plot) or the PM/vacuole fluorescence intensity ratio (dot plot) for the FAP images in panel F. Student’s t-test defines a statistically significant p-value <0.0005 in (G) and p-value <0.00005 in (H) for the WT to *pep4*Δ comparison.

While intracellular FAP-tagged probes were unaffected by the pH of the growth medium, we wondered if a cell-surface membrane protein would behave similarly. We expressed FAP-tagged Ste3 with an extracellular N-terminal FAP_optim_ tag directly exposed to environmental pH changes and quantitatively monitored its fluorescence by flow cytometry and microscopy. We observed no change in total fluorescence intensity in cells grown in a range of pHs when the fluorogen was activated using the cell permeant MG-ESTER dye. However, there was a dramatic increase in receptor internalization with decreasing pH (Figure 3E), with little-to-no receptor detected at the PM in cells that were shifted into acidic medium (pH 4.1-5.1). Notably, an acidic pH-induced increase in endocytosis has been reported previously in yeast (Motizuki *et al*., 2008). We speculate that the low pH used here caused a similar increase in endocytosis given the observed shift in FAP_optim_-Ste3 localization from the PM to intracellular compartments as the pH was reduced (Figure 3E). This pH-dependent relocalization may explain the earlier observations with FAP-Ste2 as well (Emmerstorfer-Augustin *et al*., 2018).

In contrast to the MG-ESTER results, we observed a ∼2-fold reduction in fluorescence at pH 4.1 compared to pH 7.0 with the cell impermeant MG-TAU dye (Figure 3, D and E and Supplemental Figure 2 F and G). In addition, there was a dramatic decrease in the number of cells with sufficient fluorescent signal to reach the gating threshold in flow cytometry experiments with MG-TAU at low pH (Supplemental Figure 2, F and G; <20% of cells were fluorescent). In contrast, there was no change in the percentage of cells measured across the pH ranges with the permeant MG-ESTER derivative (Supplemental Figure 2F-G). While these results might be due to pH sensitivity of the FAP/MG-TAU probe combination, given the shift in distribution of Ste3 observed with the MG-ESTER dye (Figure 3E), it more likely reflects the loss of cell surface Ste3 due to acidic pH-induced endocytosis.

During these initial experiments, we noted little FAP-tagged Ste3 vacuolar fluorescence under steady-state conditions or when the protein was internalized in response to acidic pH stress (Figure 3E). Instead, we observed several puncta peripheral to the vacuole, likely pre-vacuolar endosomes (Fig 3E-F). This result was unexpected since there is substantial Ste3 sorting to the vacuole under steady-state conditions, resulting in bright vacuolar fluorescence for receptors tagged with GFP or mCherry (Figure 3F, see Ste3-GFP) (Urbanowski and Piper, 2001; Toshima *et al*., 2014; MacDonald and Piper, 2017; MacDonald *et al*., 2020). The pH of the yeast vacuole is ∼6 (Kane, 2006), and should be compatible with robust FAP fluorescence in this compartment. However, a key feature of both GFP and mCherry is a stable beta-barrel secondary structure that is recalcitrant to digestion by vacuolar proteases and thus allows fluorescence to persist in vacuoles (Li *et al*., 1999; Nikko and Pelham, 2009).

We hypothesized that the FAP-containing SCA motif is either protease-sensitive or that FAP-tagged proteins are poorly trafficked to the vacuole. To differentiate between these scenarios, we examined FAP-Ste3 fluorescence in wildtype (WT, BY4742) cells and cells lacking the master protease Pep4, which lacks >90% of vacuolar protease activity (Woolford *et al*., 1986). Strikingly, we found that vacuolar FAP fluorescence significantly rose in *pep4*Δ cells compared to WT yeast, thereby decreasing the PM/vacuole fluorescence ratio (Figure 3F-H). Thus, the loss of vacuolar fluorescence for the FAP-tagged protein in WT cells is due to the susceptibility of the FAP tag to proteolytic digestion in the yeast vacuole and not defective trafficking of FAP-tagged proteins to the vacuole. Notably, loss of the vacuolar signal for FAP-tagged proteins is advantageous as it facilitates protein detection at other cellular locations that might otherwise be masked by bright vacuolar fluorescence, as occurs with GFP and mCherry. This vacuolar quenching makes FAP similarly useful to pHluorin, a variant of GFP that loses fluorescence in the acidic environment of the vacuole (Prosser *et al*., 2010; Prosser *et al*., 2016), but with the added adaptability of visualizing vacuolar fluorescence if needed by deleting or inhibiting vacuolar proteases (*i.e., pep4*Δ or pepstatin treatment) (Umezawa *et al*., 1970; Woolford *et al*., 1986).

### Construction of FAP_optim_ tagging vectors for yeast expression and protein co-localization studies

To provide a more widely applicable resource for the yeast cell biology community, we generated a collection of FAP_optim_-tagging vectors in which FAP-fusion protein expression can be driven from a range of promoters, each with distinct regulatory features (Figure 4A). The plasmids are based on the commonly used pRS415 or pRS413 vectors (Sikorski and Hieter, 1989; Christianson *et al*., 1992; Mumberg *et al*., 1995) that contain the *LEU2* or *HIS3* selectable markers, respectively. FAP_optim_-fusion protein expression can be optimized by incorporating the following promoters: i) *TEF1pr*, for high-level constitutive expression (Mumberg *et al*., 1995); ii) *ADH1pr*, for modest levels of constitutive expression (Mumberg *et al*., 1995); iii) *CUP1pr*, for copper-inducible expression (Labbe and Thiele, 1999); and iv) *MET25pr*, for tunable methionine-repressible expression (Kerjan *et al*., 1986) (Figure 4A). We also cloned the FAP_optim_-MYC sequence into two distinct multiple cloning site locations: i) between the *Xho*I and *Sal*I restriction sites for proteins that will be tagged at their C-terminus with FAP_optim_ or ii) between the *BamH*I and *Sma*I restriction sites for proteins that will be N-terminally tagged with FAP_optim_ (Figure 4A).

**Figure 4:**
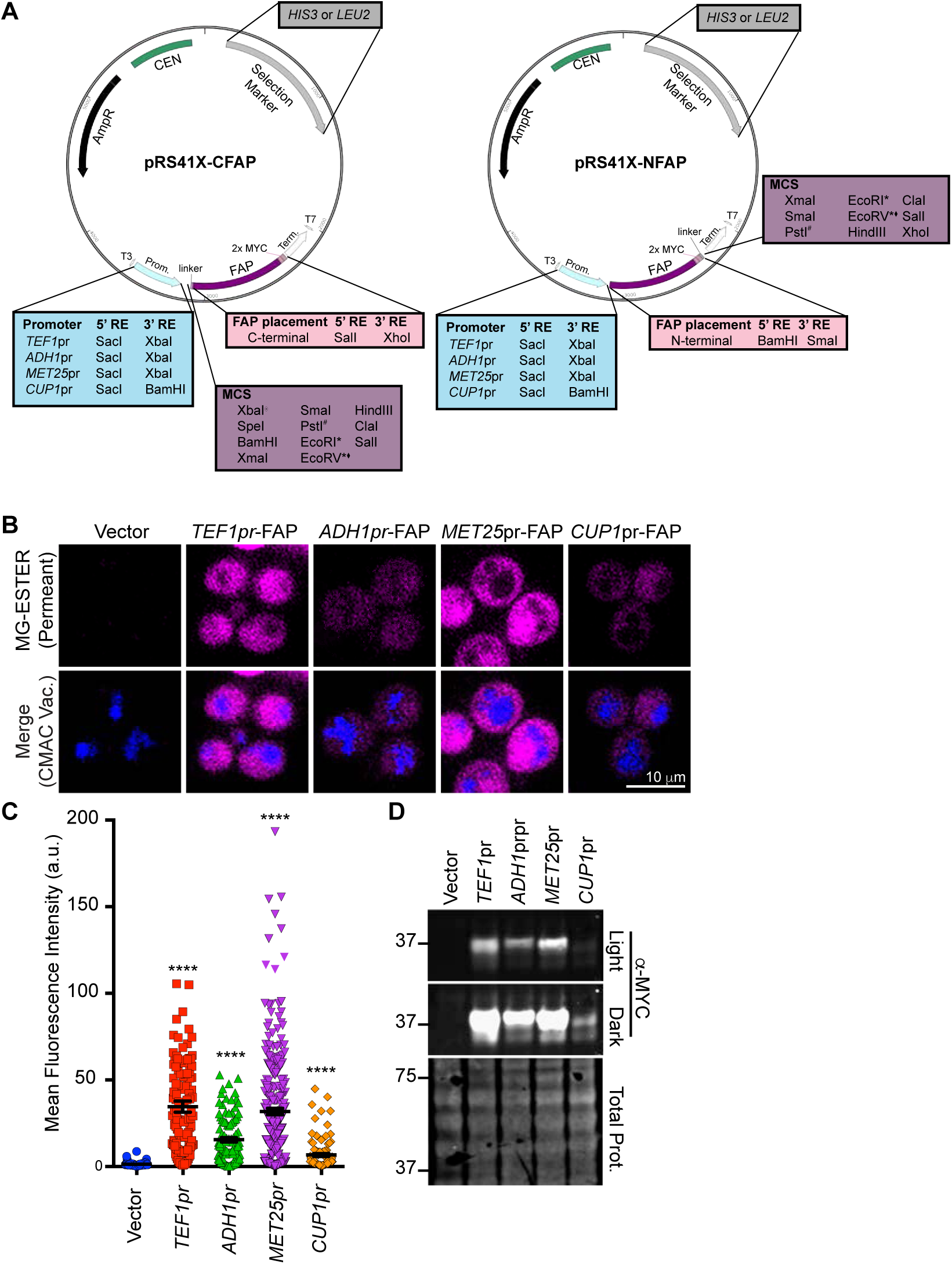
FAP-tagging vectors for use in yeast. (A) Plasmid maps of the cloning vectors built to express N- or C-terminal fusions of FAP_OPTIM_ from the promoters indicated. Plasmids contain either *HIS3* or *LEU2* for selection, the promoters and FAP sequences indicated. (B) Confocal fluorescence microscopy of soluble FAP_OPTIM_ (magenta) expressed in WT cells from plasmids containing the indicated promoters. For *MET25pr* and *CUP1pr*, images are captured at 2 h post induction. CMAC (blue) was used to visualize vacuoles. (C) Mean whole cell fluorescence intensity of FAP_OPTIM_ signal from confocal microscopy from panel B. Kruskal-Wallis with Dunn’s post hoc test was performed, and statistical comparisons are made relative to the vector only control (p<0.0005 = ***). (D) Immunoblot of whole cell extracts from WT cells expressing the FAP_OPTIM_ from the indicated promoter. For *MET25pr* and *CUP1pr*, the extracts were made from cells at 2 h post-induction. A full time-course of induction for these promoters is shown in Supplemental S3. MW markers on the left are in kilodaltons.

To assess FAP expression and ensure the promoters functioned as expected, FAP_optim_ was expressed as a soluble protein from each of the promoters in WT cells. The constitutive promoter *TEF1* and the repressible *MET25* promoter (6 h after methionine removal) yielded similar FAP-fluorescence intensities and protein expression (Figure 4, B-D). As expected, the *ADH1pr* and *CUP1pr* (120 min post copper-induction) produced significantly lower levels of FAP than the *TEF1* or *MET25* promoters (Figure 4, B-D) (Mumberg *et al*., 1995; Labbe and Thiele, 1999). An expanded time course of copper induction or methionine repression with the *CUP1* and *MET25* promoters, respectively, to determine the optimal induction timing for FAP_optim_, is shown in Supplemental Figure 3A-F. We also inserted an ER-targeting sequence upstream of the N-terminal FAP_optim_ tag to facilitate protein entry into the secretory pathway (Figure S4A). Finally, we modified constructs to facilitate chromosomal integration of the FAP tag (Figure S4B). In sum, we generated 17 new cloning vectors for expressing or tagging FAP proteins (Table S2) to maximize the system’s versatility.

To further aid in the implementation of the FAP technology in yeast research, we created a suite of cellular proteins tagged with FAP_optim_ for use in co-localization studies (Table S2). These probes were cloned into the pRS413-*TEF1pr* plasmid for constitutive expression (Figure 5A; Table S2). We created FAP_optim_-tagged markers for the *cis*-Golgi (Anp1), lipid droplets (Erg6), the PM (Pma1), the nucleus (Rpa34), the *trans*-Golgi (Sec7), the ER (Sec61 and Sec63), and the vacuolar membrane (Vph1) (Deshaies and Schekman, 1987; Kane, 1999; Todorow *et al*., 2000; Young *et al*., 2001; Glick and Nakano, 2009; Albert *et al*., 2011; Young *et al*., 2024). To ensure correct localization, markers were co-expressed in cells in which the respective endogenous protein was tagged with either RFP (Huh *et al*., 2003) or mNeonGreen (mNG) (Meurer *et al*., 2018), and co-localization was assessed by live cell fluorescence microscopy in the presence of MG-ESTER (Figure 5B-C). For all the organelle markers examined, we confirmed that the FP-and FAP-tagged proteins colocalized at the expected locations in the cell (Figures 5B-C). While these findings show that the FAP-and FP-tagged organelle marks behave similarly, it should be noted that we did not functionally assess these FAP-tagged probes (*i.e.,* functional complementation assays have not been done for all FAP-tagged organelle markers).

**Figure 5:**
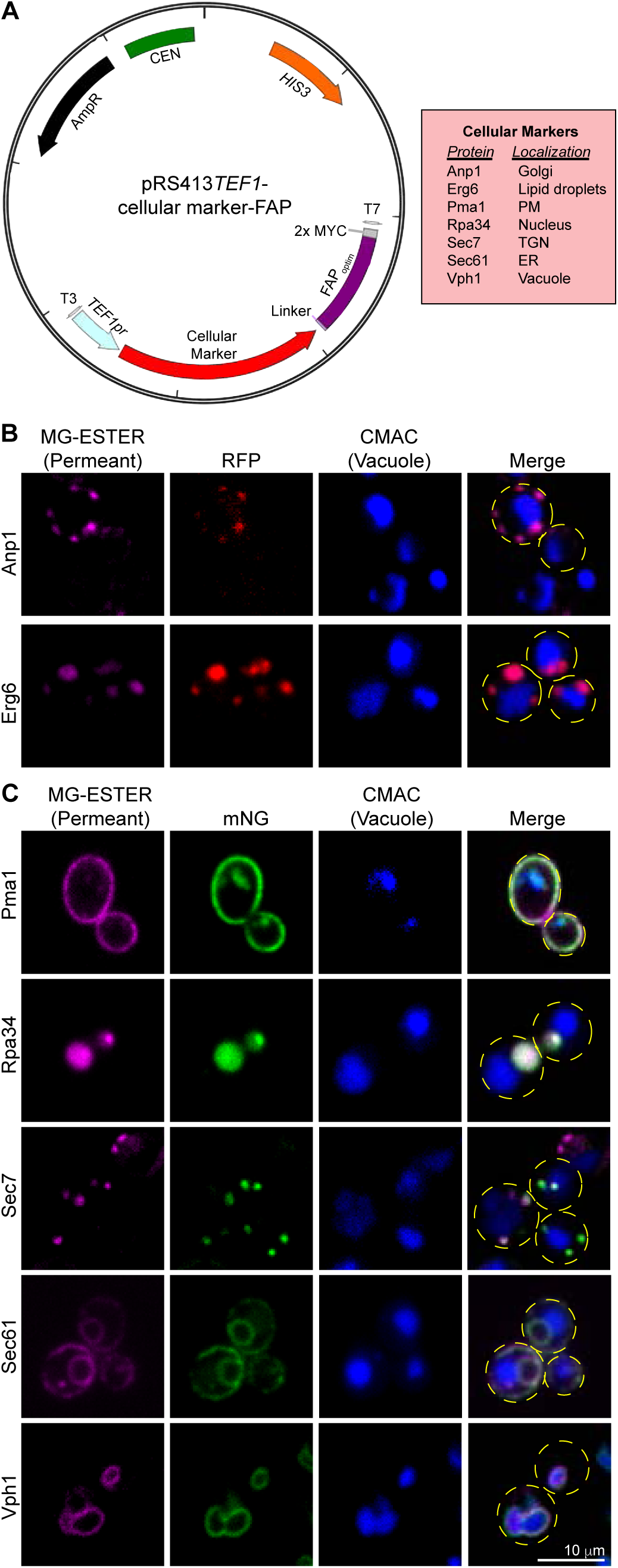
FAP-tagged cellular markers. (A) Schematic of plasmids expressing C-terminal fusions to FAP_OPTIM_ for the genes indicated in the red box at right. (B-C) Confocal fluorescence microscopy of WT cells expressing the FAP_OPTIM_-tagged proteins indicated (magenta) as well as the same protein RFP-tagged (panel B, red) or mNG-tagged (panel C, green). Cell outlines are shown as yellow dashed lines in the merge for reference, and CMAC (blue) stains the vacuoles.

### FAP_optim_-tagged Ste3 is a functional a-factor receptor

We next sought to use the FAP_optim_ system to define the contributions of *cis*- and *trans*-acting regulators of ligand-induced Ste3 trafficking. The dynamics of Ste3 post pheromone (a-factor) addition have not yet been characterized by live cell imaging as previous studies relied predominantly on biochemical assays to define the Ste3 trafficking itinerary (Zanolari *et al*., 1992; Chen and Davis, 2000, 2002). These past studies identified regulatory sequences within the C-tail of Ste3 that control its internalization. Trans-acting regulators have also been described, mainly for the constitutive endocytosis and recycling of Ste3 (Chen and Davis, 2002; Prosser *et al*., 2015; MacDonald and Piper, 2017; Laidlaw *et al*., 2022a; Laidlaw *et al*., 2022b). Here, we use FAP-tagged Ste3 to assess the role of these regulators in ligand-induced endocytosis of the receptor. Previously, FP-based studies of Ste3 endocytic dynamics were thwarted by the fact that bright fluorescent signal accumulates in the vacuoles from FP-tagged Ste3 (Urbanowski and Piper, 2001; MacDonald and Piper, 2017), which occluded detection of tagged Ste3 at other intracellular locales. In addition, Ste3 gene expression is dramatically increased post a-factor addition (Sprague *et al*., 1983), muddying isolated analysis of the pre- and post-ligand expressed pools. However, FAP-Ste3 allows us to fluorescently label a single pool of this receptor at the PM and monitor its localization over time in a fluorescence-based ‘pulse-chase’ assay.

We first confirmed that the FAP-Ste3 protein exhibits WT activity, akin to untagged, endogenous Ste3, or the extensively studied Ste3-GFP fusion protein (Urbanowski and Piper, 2001; MacDonald and Piper, 2017; MacDonald *et al*., 2020). Upon activation of the mitogen-activated protein kinase (MAPK) cascade downstream of Ste3, yeast cells undergo a morphological shift known as ‘shmoo’ formation (Bardwell *et al*., 1994; Bardwell, 2005), which facilitates the fusion of mating cells. When expressed as the sole copy of Ste3, we found that FAP-Ste3 was as efficient at stimulating shmoo formation as endogenous Ste3 or Ste3-GFP (Figure 6A-B). In contrast, *ste3*Δ cells expressing FAP_optim_ lacked shmoos (Figure 6A-B). Next, because the addition of the Ste3 ligand, a-factor, induces expression of the MAPK, Fus3 (Couve and Hirsch, 1996), we assessed Fus3 protein levels. We found that Fus3 increased in a-factor treated cells expressing FAP-Ste3 with similar kinetics to those observed for WT cells with an empty vector or *ste3*Δ cells expressing Ste3-GFP (Figure 6C). Finally, we used halo assays to assess the ability of FAP-Ste3 to induce the hallmark growth arrest associated with activating the mating pathway in response to a-factor. Cells expressing endogenous, untagged Ste3, or *ste3*Δ cells expressing FAP-Ste3 or Ste3-GFP produced halos of equivalent sizes (Figure 6 D-E). Thus, FAP-Ste3 is a functional, a-factor-stimulated GPCR suitable for studies to monitor the mechanism of Ste3 trafficking.

**Figure 6:**
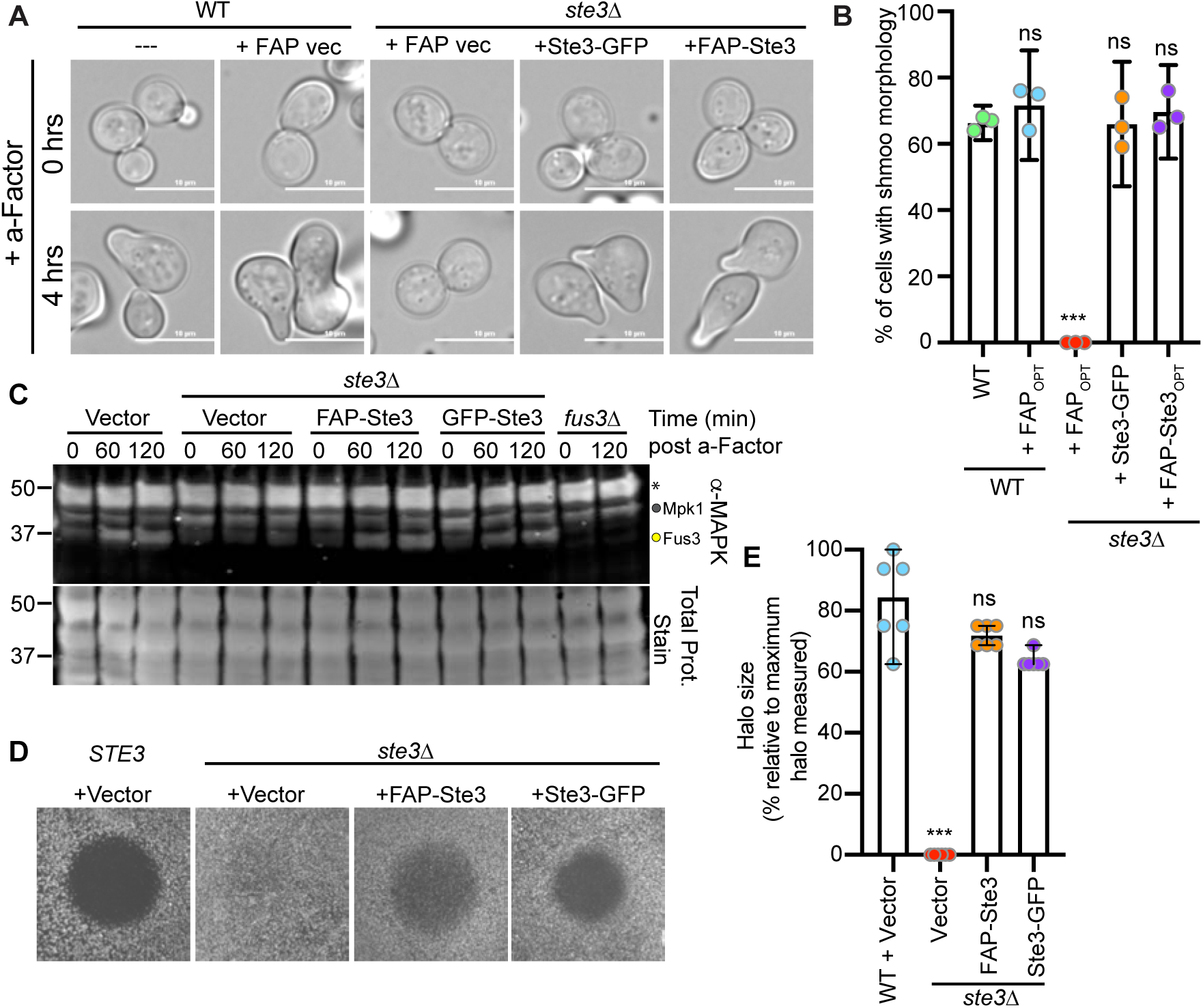
FAP-tagged Ste3 is a functional GPCR in the mating pathway. (A) FAP_optim_, Ste3-GFP, or FAP_optim_-Ste3 were expressed from the *TEF1pr* in WT or *ste3*Δ cells and imaged by microscopy before and after 4 h treatment with 5 μM a-factor. (B) Quantification of the percentage of cells with a shmoo morphology (three biological replicates) from the imaging in panel (A). Mean with 95% confidence interval shown. Mann-Whitney t-test was performed comparing each set to WT (not significant = ns; *p*-value < 0.001 = ***). (C) Immunoblot of protein extracts from cells treated with 5 μM a- factor for the indicated times. The band indicated with the yellow circle is Fus3, the MAPK activated by the mating pathway; the band immediately above (grey circle) is Mpk1, another MAPK in yeast that is also detected with this antibody; and the top band (asterisks) is an unknown cross-reacting band. (D) Representative a-factor halo assays, where 5 μg a-factor was spotted at the center of these halos (filter removed for visualization) for either WT or *ste3*Δ cells expressing the indicated plasmids. (E) Quantification of halo diameters as shown in panel D represented as a percentage of the maximal halo size measured across 6 biological replicates. Student’s t-test compared experimental values to WT + vector (not significant = ns; p<0.0005 = ***)

### Deletion of yapsin Mkc7 improves PM detection of FAP-Ste3

An earlier study using FAP-tagged Ste2, which resides at the PM of *MATa* cells, found that SCA-dependent fluorescence and receptor stability were improved when the yapsins—glycosylphosphatidylinositol-anchored aspartyl proteases on the cell surface (Krysan *et al*., 2005)—were deleted (Emmerstorfer-Augustin *et al*., 2018). This increase was likely due to loss of yapsin-dependent cleavage of the FAP tag from Ste2. Consistent with this prior study, we found that deletion of the yapsin Mkc7 significantly increased surface FAP-Ste3 fluorescence in the presence of the cell-impermeant MG-TAU dye (Figure 7A-B); the absence of another yapsin, Yps1, did not further increase surface fluorescence (Figure 7A-B). Deleting these aspartyl proteases facilitated the quantitative detection of steady-state and ligand-induced FAP-Ste3 internalization (Figure 7C-F). Importantly, a-factor induced internalization of FAP-Ste3 was faster (complete after ∼20 min) than the observed constitutive internalization of Ste3, in which internalization continued for ∼60 min (compare Figures 7C-D to 7E-F) in the absence of yapsins. We note, however, that Ste3 internalization was challenging to observe in cells where the yapsins were intact (*i.e.,* WT cells) because the initial fluorescent signal for FAP-Ste3 was low, and possibly confounded by ongoing FAP cleavage (Figures 7C-F).

**Figure 7:**
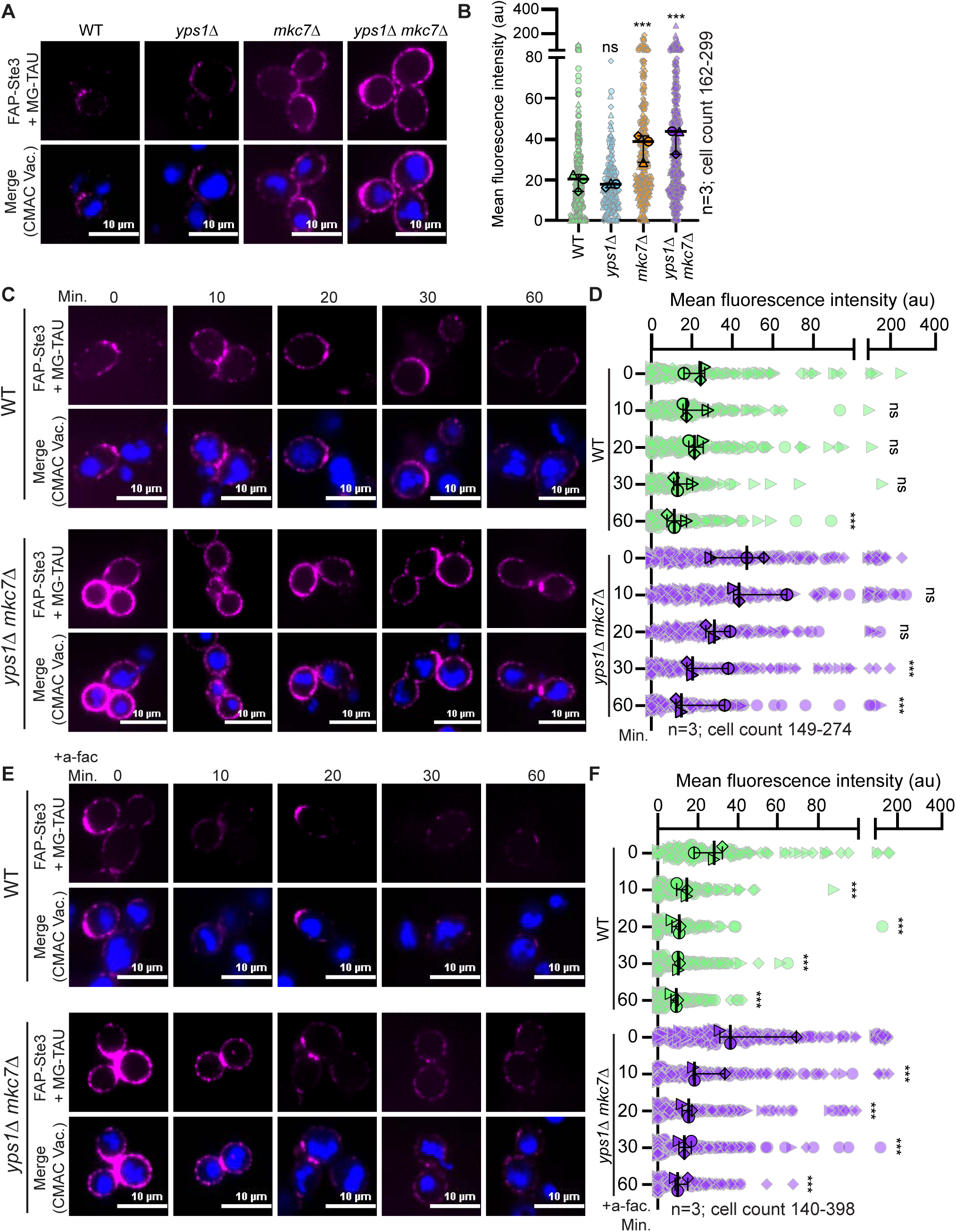
Deletion of yapsin Mkc7 improves FAP-Ste3 fluorescence. (A) Representative confocal microscopy images of FAP_optim_-Ste3 expressed from the *STE3pr* in the genetic backgrounds indicated. Cells were incubated with MG-TAU (impermeant) dye to visualize the surface pool of Ste3 (magenta) and CMAC (blue) stained the vacuoles. (B) Whole-cell fluorescence intensities from cells imaged in panel A were determined using ImageJ for three biological replicates. Data are presented as a Superplot where each cell measured is plotted as a grey outlined shape (circle, triangle, diamond), with the shapes indicating cells from one of the 3 replicates. The mean fluorescence intensity of each replicate is then plotted as a larger shape outlined in black with the mean for each of the three trials and 95% confidence interval shown with black bars. Kruskal-Wallis statistical analysis with Dunn’s post hoc test was performed to compare the means of the three replicates to the WT control (not significant = ns; p<0.0005 = ***). (C & E) Representative confocal fluorescence microscopy images of FAP_optim_-Ste3 expressed from the *STE3pr* in WT and *yps1*Δ *mkc7*Δ cells either untreated (C) or treated with a-factor (E). Cells were incubated with MG-TAU (impermeant) dye at t=0 to visualize cell surface Ste3 and CMAC-stained vacuoles. The dye was then washed from the cells, and cells were imaged over time (C) or 5 μM a-factor was added, and cells were imaged over time (E). This allowed us to monitor the steady-state (panel C) or ligand-induced (panel E) turnover of FAP-Ste3 from the PM. (D & F) Whole-cell fluorescence intensity for cells imaged as in panels (C) or (E), respectively, was determined using ImageJ for three biological replicate experiments. The data are presented as a Superplot where each cell measured is plotted as a grey outlined shape, with the shapes corresponding to a single replicate. The mean fluorescence intensity for each replicate is plotted as a circle, triangle, or diamond outlined in black with the mean for each of the three trials and 95% confidence interval shown with black bars. Kruskal-Wallis statistical analysis with Dunn’s post hoc test was performed to compare the means of the three replicates to the WT control (not significant = ns; p<0.0005 = ***).

### Assessment of cis-acting sequences on ligand-induced Ste3 endocytosis

The posttranslational modification of the C-terminal tail of many GPCRs regulates endocytosis and trafficking by mediating key interactions with trafficking regulators (Roth and Davis 2000, Zanolari et al 1992, Dohlman and Thorner 2001). Like other GPCRs, Ste3 has a long C-terminal tail implicated in its endocytosis (Zanolari and Riezman, 1991; Chen and Davis, 2000, 2002). Next, we evaluated the role of Ste3 C-terminal sequences in regulating ligand-induced turnover. Prior studies of these amino acid motifs and their role in Ste3 internalization were primarily limited to biochemical assessment (Roth and Davis, 1996; Roth *et al*., 1998; Roth and Davis, 2000), as quantitative assessment of Ste3-GFP endocytosis by live-cell imaging was confounded by high levels of vacuolar Ste3-GFP fluorescence. We generated a FAP-Ste3-Tailless mutant lacking the entire C-terminal tail (amino acids 288-470) and monitored its turnover from the PM relative to the FAP-Ste3 WT control. We found that a-factor-induced turnover of Ste3-Tailless was significantly disrupted, resulting in prolonged receptor retention at the cell surface (Figure 8A-B). Ste3-Tailless remained at the PM for up to 30 minutes post a-factor addition, whereas nearly all WT receptors were internalized in this timeframe (Figure 8A-B).

**Figure 8:**
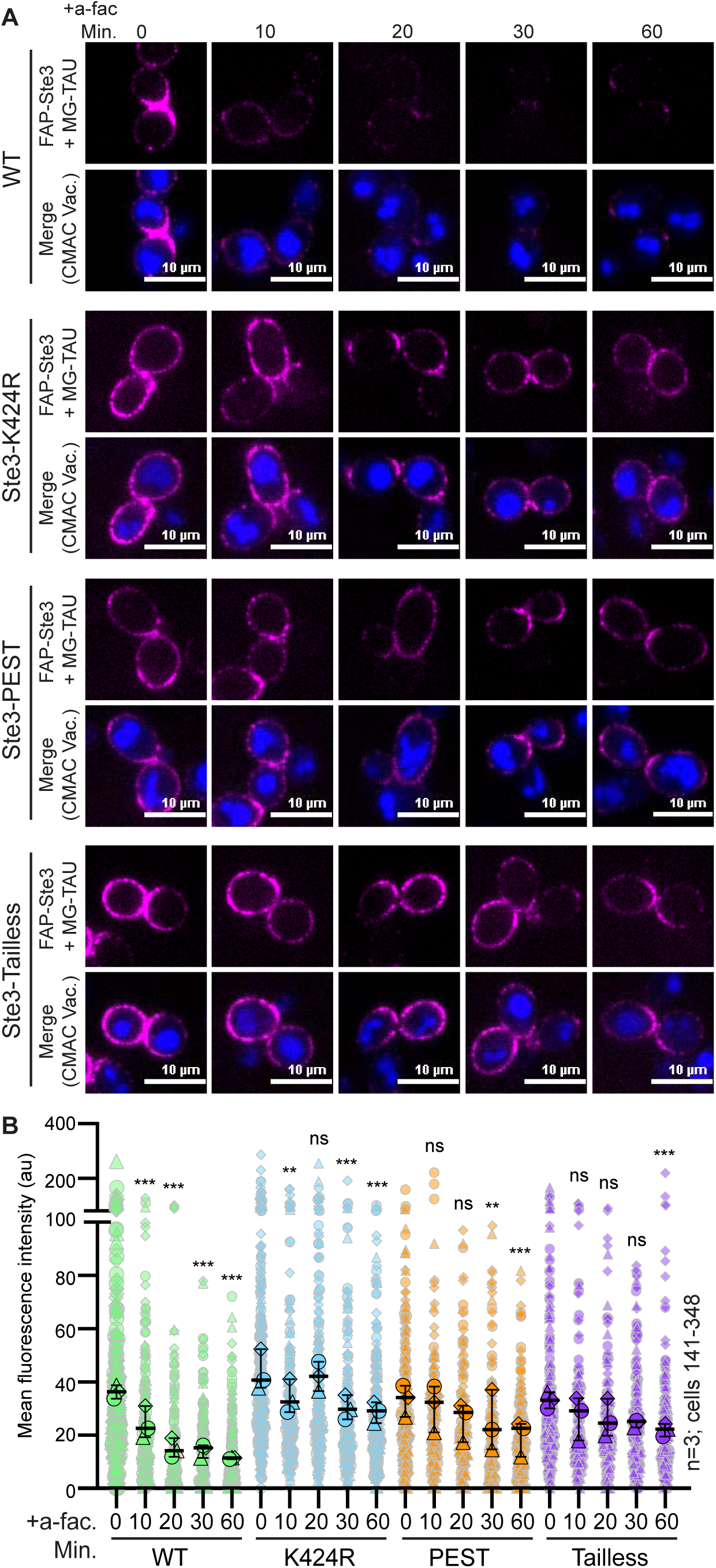
The impact of *cis*-acting sequences on ligand-induced turnover of FAP-Ste3. (A) Representative confocal fluorescence microscopy images of either WT or mutated FAP_optim_-Ste3 expressed from the endogenous *STE3pr* in *yps1*Δ *mkc7*Δ cells treated with 5 μM a-factor for the indicated time. Cells were incubated with MG-TAU (impermeant; magenta) dye at t=0 to visualize cell surface Ste3 and CMAC (blue) stained the vacuoles. The dye was washed from the cells, 5 μM a-factor was added, and cells were imaged over time, allowing ligand-induced turnover of FAP-Ste3 to be monitored. (B) Whole-cell fluorescence intensity for cells imaged in panel A was determined using ImageJ for three biological replicate experiments. The data are presented as a Superplot where each cell measured is plotted as a grey outlined shape, with the shapes corresponding to a single replicate. The mean fluorescence intensity for each replicate is plotted as a larger circle, triangle, or diamond outlined in black with the mean for each of the three trials and 95% confidence interval shown with black bars. Kruskal-Wallis statistical analysis with Dunn’s post hoc test was performed to compare the means of the three replicates to t=0 control for each version of Ste3 (not significant = ns; p=<0.005 = **; p<0.0005 = ***).

To further refine our analysis of the Ste3 C-tail, we deleted the PEST-like motif (Ste3-ΔPEST; amino acids 413-470), posited to regulate constitutive, but not ligand-induced, Ste3 endocytosis (Roth *et al*., 1998). While the defect in ligand-induced turnover of Ste3-ΔPEST was more modest than that observed for Ste3-Tailless, it was significantly slowed compared to WT Ste3 (Figure 8A-B). These findings suggest that the PEST sequence not only regulates constitutive turnover but also controls ligand-induced Ste3 internalization.

Finally, we assessed the role of ubiquitination in ligand-induced Ste3 internalization. Selective ubiquitination of membrane proteins at key cytosolic lysine residues is often a critical signal for their endocytosis (Rotin *et al*., 2000; Herrador *et al*., 2013), and early studies of Ste3 demonstrate ubiquitination of the Ste3 C-terminal tail regulates turnover, at least in biochemical assays (Roth and Davis, 1996; Roth *et al*., 1998; Roth and Davis, 2000). Though the entire C-tail of Ste3 contains 17 lysines that could serve as putative ubiquitination sites, only three lysines residing within the PEST domain have been shown to alter Ste3 ubiquitination and internalization (Roth and Davis, 1996; Roth *et al*., 1998; Roth and Davis, 2000). While past studies indicate that these three lysines (K424, K432, and K453) are somewhat functionally redundant, mutation of K424 to arginine alone significantly delayed Ste3 turnover (Roth and Davis, 2000). Therefore, we made the FAP-tagged Ste3-K424R mutant and assessed endocytic turnover in response to a-factor. Like the PEST deletion, this single point mutation impaired loss of FAP-Ste3 signal at the PM post ligand addition, consistent with reduced endocytic turnover (Figure 8A-B). Consistent with the importance of K424 ubiquitination, it was difficult to distinguish a difference between Ste3-Tailless, -K424R, or Ste3-ΔPEST kinetics (Figure 8A-B).

### Ligand-Induced endocytosis of Ste3 is regulated by the Aly1, Aly2 and Art1 α-arrestins

We previously reported that the constitutive endocytosis of Ste3-pHluorin depends on α-arrestins (Prosser *et al*., 2015), a class of protein trafficking adaptors conserved from yeast to man (Alvarez, 2008; Lin *et al*., 2008; Nikko and Pelham, 2009; O’Donnell *et al*., 2010; O’Donnell and Schmidt, 2019). α-Arrestins serve as a bridge between membrane proteins and the Rsp5 ubiquitin ligase, which ubiquitinates membrane proteins to permit their efficient endocytosis (Lin *et al*., 2008; Nikko and Pelham, 2009; Becuwe *et al*., 2012; O’Donnell *et al*., 2013). In our past work, of the 14 yeast α-arrestins, only Aly1, its paralog Aly2, and Art1 (aka Art6, Art3, and Ldb19, respectively) stimulated Rsp5-dependent internalization of Ste3 under steady-state conditions (Prosser *et al*., 2015). Importantly, these findings are supported by analogous studies of FAP-Ste3 (Supplemental Figure S5A-B). By examining FAP-Ste3 in combination with the MG-TAU dye, we selectively monitored Ste3 abundance at the PM after a-factor addition. As observed when constitutive internalization was assessed (Prosser *et al*., 2015), cells lacking either Art1, Aly1 and Aly2, or all three of these α-arrestins, exhibit delayed Ste3 internalization after a-factor addition (Figure 9A-B). Emphasizing the utility of the FAP- tagging system, when we similarly monitored ligand-induced endocytosis using Ste3-pHluorin in WT cells or those lacking α-arrestins, no significant drop in Ste3 fluorescence was observed (Figure 10 A-B and Supplemental Figure S6A-B).

**Figure 9:**
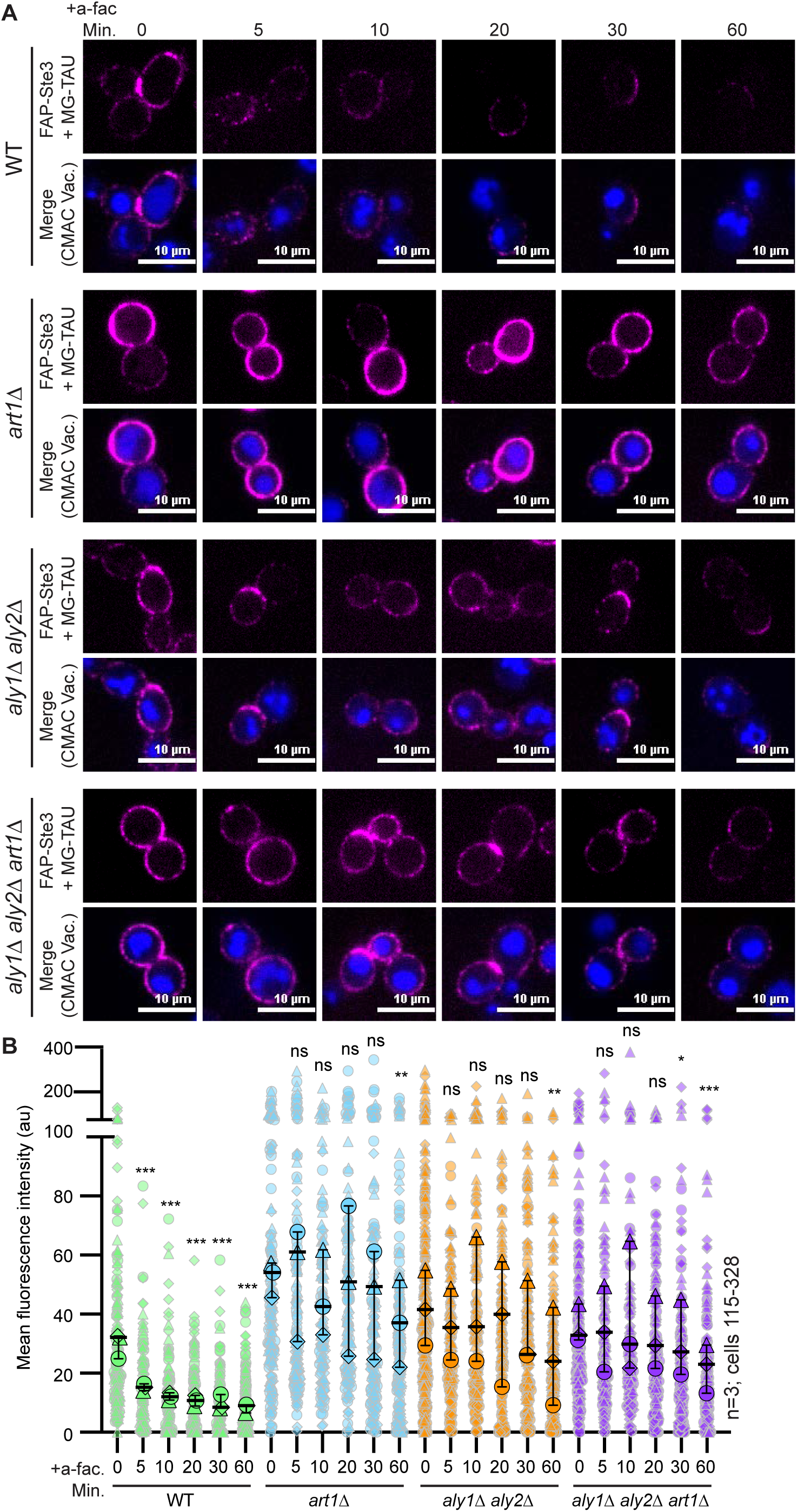
Contribution of α-arrestins to ligand-induced turnover of FAP-Ste3. (A) Representative confocal fluorescence microscopy images of FAP_optim_-Ste3 expressed from the endogenous *STE3pr* in the cells indicated and treated with 5 μM a-factor. Cells were incubated with MG-TAU (impermeant) dye at t=0 to visualize cell surface Ste3, and CMAC stained the vacuoles. The dye was then washed from the cells, 5 μM a-factor was added, and cells were imaged over time. This allows us to monitor the ligand-induced turnover of FAP-Ste3 from the PM. (B) Whole-cell fluorescence intensity for cells imaged in panel A was determined using ImageJ for three biological replicate experiments. The data are presented as a Superplot where each cell measured is plotted as a grey outlined shape, with the shapes corresponding to a single replicate. The mean fluorescence intensity for each replicate is plotted as a circle, triangle, or diamond outlined in black with the mean for each of the three trials and 95% confidence interval shown with black bars. Kruskal-Wallis statistical analysis with Dunn’s post hoc test was performed to compare the means of the three replicates to t=0 control for each of the strains (not significant = ns; p=<0.05 = *; p=<0.005 = **; p<0.0005 = ***).

**Figure 10:**
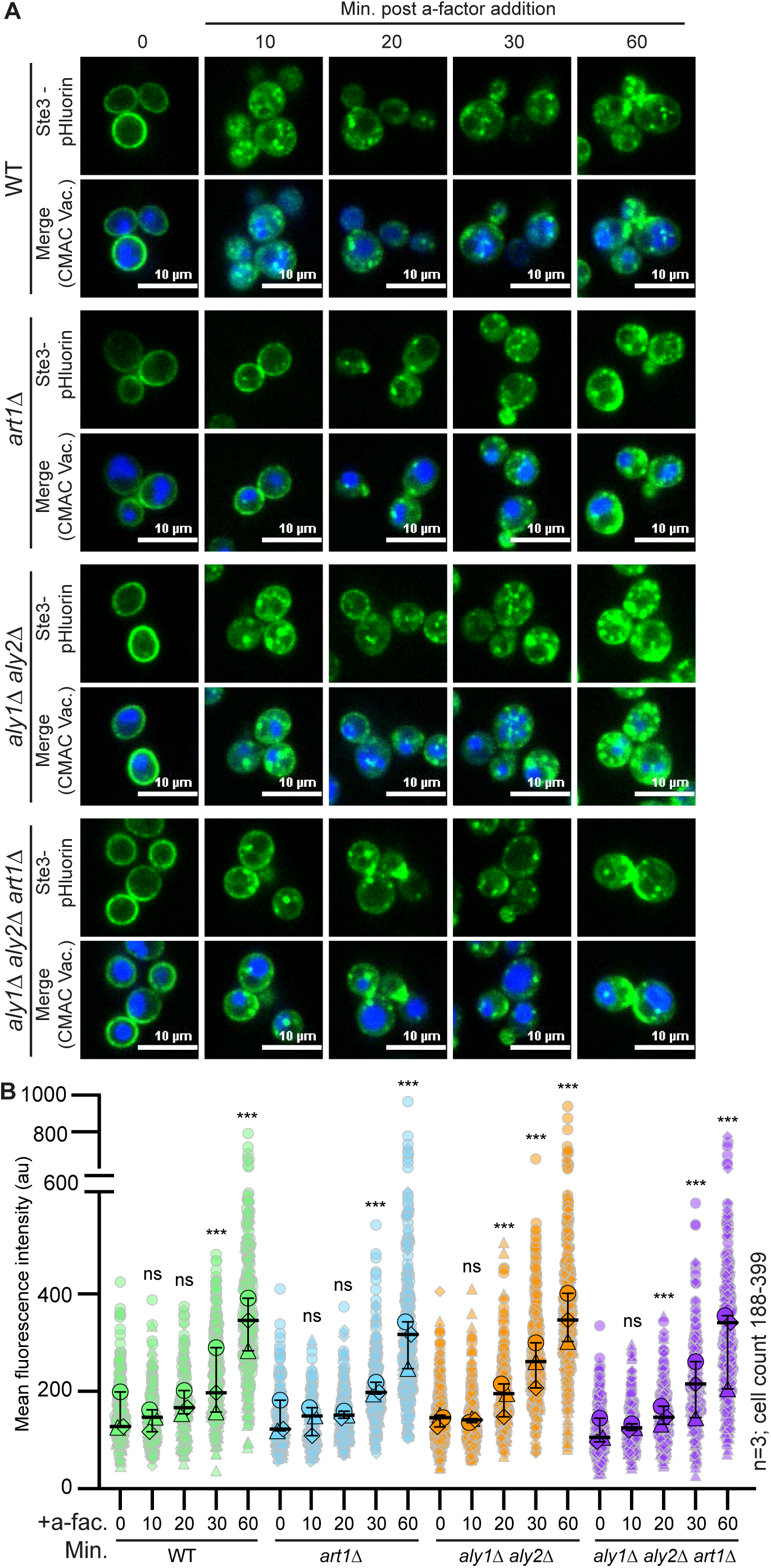
Using pHluorin-Ste3 to monitor ligand-induced turnover of the receptor. (A) Representative confocal fluorescence microscopy images are presented of Ste3-pHluorin (green) expressed from the endogenous *STE3* chromosomal locus in either WT or cells lacking the indicated α-arrestin. Cells were either untreated or treated with 5 μM a-factor and CMAC (blue) stained the vacuoles. (B) Whole-cell fluorescence intensity for cells imaged in panel A was determined using ImageJ for three biological replicate experiments. The data are presented as a Superplot where each cell measured is plotted as a grey outlined shape, with the shapes corresponding to a single replicate. The mean fluorescence intensity for each replicate is plotted as a circle, triangle, or diamond outlined in black with the mean for each of the three trials and 95% confidence interval shown with black bars. Kruskal-Wallis statistical analysis with Dunn’s post hoc test was performed to compare the means of the three replicates to t=0 control for each strain (not significant = ns; p<0.0005 = ***).

To ensure that Ste3-pHluorin and FAP-Ste3 behaved similarly, we monitored total FAP-Ste3 post a-factor addition by adding the MG-ESTER dye at every interval during a time course. This contrasts with our prior experiments where we labelled FAP-Ste3 with MG-TAU at the zero timepoint and then monitored the change in distribution for this subset of the receptor (Figure 8A-B). When MG-ESTER dye is added at each timepoint, we should observe the entire FAP-Ste3 population, which includes the intracellular and cell surface pools of Ste3 prior to ligand induction and newly transcribed and translated Ste3 after the ligand is added. By adding MG-ESTER dye over time, we found that, like Ste3-pHluorin, FAP-Ste3 also increases in abundance after a-factor addition, and these two differentially tagged receptors co-localized (Figure 11A-B). Thus, pHluorin- and FAP-tagged Ste3 behave similarly in our assays when the total pool of each protein is monitored.

**Figure 11:**
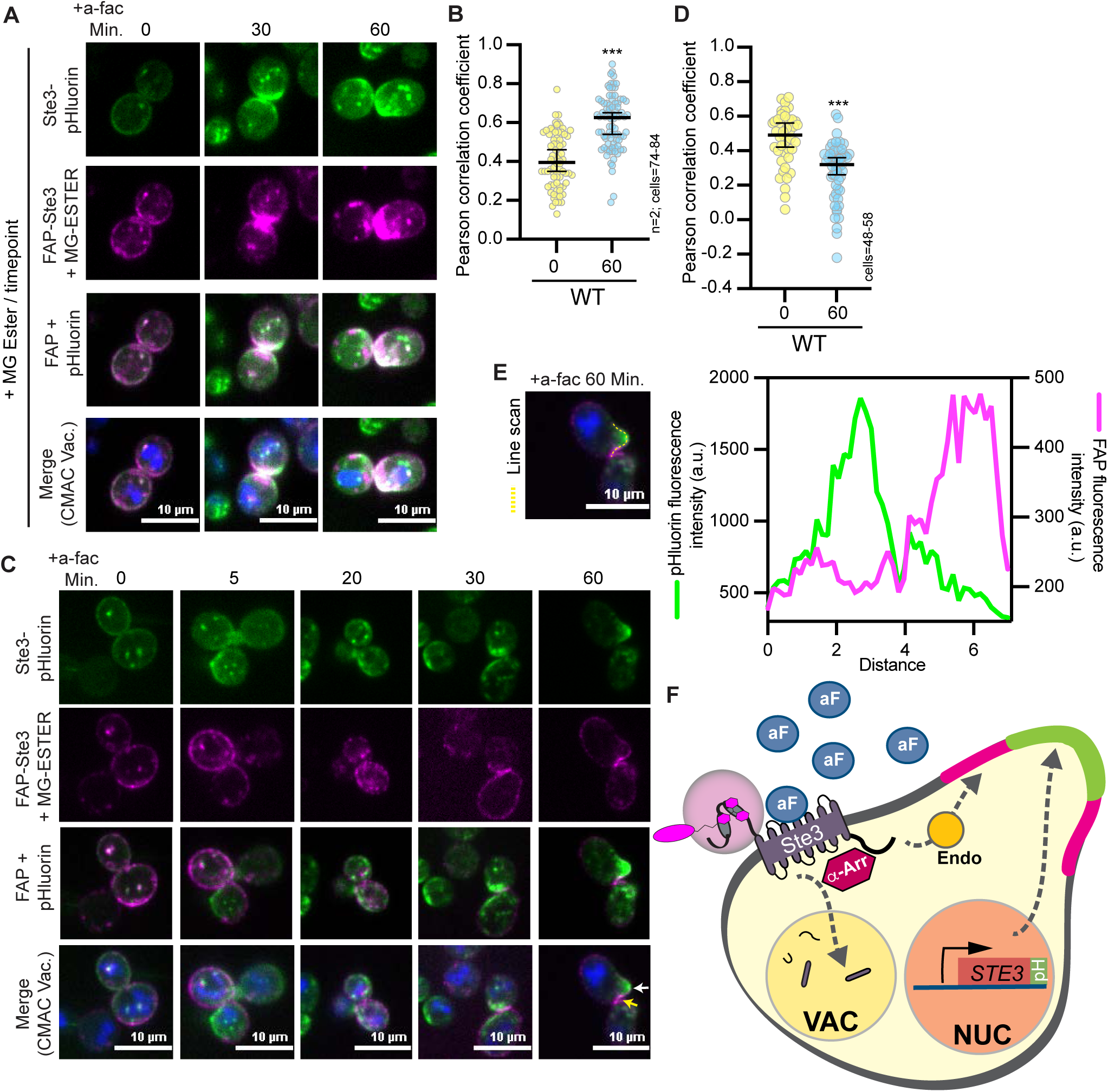
FAP-Ste3 can be used to monitor recycling from the PM. (A) Cells expressing FAP_optim_-Ste3 from a plasmid using the *STE3pr* and chromosomally integrated Ste3-pHluorin were imaged by confocal fluorescence microscopy. Cells were incubated with MG-ESTER and CMAC dye. Once the dye was washed from the cells, 5 μM a-factor was added and the cells were imaged at the indicated timepoints <u>with fresh</u> <u>MG-ESTER dye added prior to imaging each timepoint</u>. (B and D) Pearson’s correlation coefficient of the FAP-Ste3 and Ste3-pHluorin fluorescent signals in panels A and C, respectively, were determined using the colocalization analysis plugin in Image J. (C) Cells expressing FAP_optim_-Ste3 from a plasmid using the *STE3pr* and chromosomally integrated Ste3-pHluorin were imaged by confocal fluorescence microscopy. Cells were incubated with MG-ESTER and CMAC dye. Once the dye was washed from the cells, 5 μM of a-factor was added, and the cells were imaged at the indicated timepoints with no further dye additions. This allowed us to monitor a single pool of Ste3 at the onset of the experiment with FAP while monitoring the total pool of Ste3 with pHluorin. The yellow arrow points to FAP-Ste3 at the side of the shmoo while the white arrow points to Ste3-pHluorin at the tip of the shmoo. (E) Line scan analysis to monitor the relative distributions of FAP-Ste3 and Ste3-pHluorin at the shmoo tip after pheromone addition. The region used in the line scan is indicated as a dashed yellow line on the image and the fluorescence intensities along this line for GFP (left y-axis) or FAP (right y-axis) are plotted. (F) Model of Ste3 trafficking based on FAP and pHluorin-tagged studies. The addition of Ste3’s ligand, a-factor (aF, blue circle), induces MAPK signaling from the Ste3 receptor and leads to morphology and transcriptional changes in cells that allow for mating. As part of the transcriptional response, *STE3* gene expression is induced, generating new Ste3 protein that is pHluorin tagged. Newly synthesized FAP-Ste3 will not be visualized unless the dye is added at each timepoint (as in Fig 11A). The FAP-Ste3 pool present at the PM prior to a-factor addition can be visualized selectively using the cell impermeant dye administered just before a-factor treatment (as in Fig 11C). Newly synthesized Ste3-pHluorin localizes to the shmoo tip of cells while “old” FAP-Ste3 that was at the PM prior to a-factor addition localizes to the sides of the shmoo tip. It is not clear which pathways control these distinct targeting events for Ste3. We find that α-arrestins (Aly1, Aly2 and Art1) are important for the ligand-induced internalization of Ste3 and that the C-tail of Ste3 is needed for its efficient endocytosis. Past work shows that the C-tail is also the site of α-arrestin binding, and the PEST and K424 residues are important cis-regulators of Ste3 internalization. Nuc=nucleus; Vac=vacuole; Endo=endosome.

If we further alter the parameters for the imaging assay, and instead only add MG-ESTER at a single time point prior to a-factor addition, we can selectively monitor the pool of FAP-Ste3 that is expressed prior to a-factor addition. In contrast, the Ste3-pHluorin reports on both the pre-existing pool of Ste3 and the newly synthesized Ste3 post a-factor addition. In this assay, FAP-Ste3 and Ste3-pHluorin initially colocalized (Figure 11C, timepoints 0, 5 and 10 min and Figure 11D), but after prolonged incubations with a-factor, the pre-existing FAP-labelled pool differed relative to the pool of Ste3-pHluorin (Figure 11C-D, 60 min timepoint). Specifically, by 1 h post a-factor addition, the Ste3-pHluorin pool accumulated at the shmoo tip, whereas FAP-Ste3, which represents the preexisting pool of Ste3 before a-factor addition, partitioned to the side of the shmoo (Figure 11C and E and Supplemental Figure S6A-B). These findings suggest differential targeting of the preexisting receptor—present before ligand induction and represented by the FAP-Ste3 population at the sides of the shmoo tip— versus newly synthesized and exocytosed receptor, which is expressed after a-factor addition and is defined by the Ste3-pHluorin population at the shmoo tip (Figure 11F). In the future, it will be exciting to use these new FAP tools to better define the features that dictate this differential sorting of “old” and “new” Ste3 receptor populations.

## DISCUSSION

### Adaptation of FAP for imaging in yeast

Current approaches to studying endocytosis and intracellular trafficking of membrane proteins are limited by their inability to monitor the dynamics of select protein sub-populations. To overcome this challenge, we developed an optimized FAP live-cell imaging system for use in *S. cerevisiae* (Perkins and Bruchez, 2020). To facilitate the use of FAP in yeast imaging studies, we constructed plasmids for building FAP fusions or making FAP-tagged integrations, as well as a suite of FAP-tagged co-localization markers. In our current work, FAP is imaged in the far-red channel (640 excitation/680 emission) when bound to the cell-permeant MG-ESTER dye, allowing for the total pool of protein to be monitored, or the cell-impermeant MG-TAU dye, permitting the surface population of a protein to be analyzed in isolation (Figure 1). These features make FAP a highly useful probe as few current live cell fluorescent tags exploit the far-red channel, which is spectrally distinct from commonly used green, red, and blue fluorophores (*i.e.,* GFP, RFP, BFP and their derivatives). Importantly, most other approaches fail to differentially track specific protein subpopulations, as can be achieved with this dual FAP dye system.

To improve the parameters for FAP imaging in yeast, we i) altered the nucleotide sequence encoding FAP to ensure that the optimal codons for expression and stability in yeast were used, ii) identified the pH range (4.1-7) over which FAP fluorescence is detectable, iii) demonstrated that low pH (<6.0) can induce endocytosis of membrane proteins and should be avoided for trafficking studies in yeast, iv) found that the FAP tag is susceptible to cleavage by vacuolar proteases, which greatly diminishes FAP fluorescence when the protein is sorted to this organelle, and v) confirmed that when FAP is expressed extracellularly it is cleaved by the yapsin aspartyl proteases, demonstrating that, for quantitative endocytic studies, the yapsins should be deleted from the cell. With these parameters defined, we used FAP technology to study ligand-induced endocytosis of the Ste3 GPCR.

### Ligand-induced Ste3 endocytosis and recycling

The function of Ste3 and its downstream signaling pathway in controlling yeast mating have been actively investigated for decades (Bardwell *et al*., 1994; Bardwell, 2005). Initially, biochemical studies defined the machinery that mediates Ste3 endocytosis and post-endocytic recycling of Ste3 to the PM (Roth and Davis, 1996; Roth *et al*., 1998; Chen and Davis, 2000; Roth and Davis, 2000; Chen and Davis, 2002). Importantly, many cell surface receptors in mammals analogously recycle to the PM after internalization, and some are dually targeted for post-endocytic recycling to the PM and degradation in the lysosome (Tomas *et al*., 2014). While the endocytosis of many membrane proteins is ubiquitin-dependent, post-endocytic sorting to the vacuole is far better described than the endosomal recycling pathways, which are still being characterized (Bonifacino and Weissman, 1998; Sorkin and Von Zastrow, 2002). For Ste2, the counterpart of Ste3 in *MATa* cells, endocytosis leads to vacuolar targeting and degradation (Schandel and Jenness, 1994; Urbanowski and Piper, 2001; MacDonald and Piper, 2017; MacDonald *et al*., 2020). However, protease-shaving assays measuring Ste3 PM abundance after a-factor treatment suggested that internalized Ste3 is recycled to the PM (Chen and Davis, 2000). Ste3 recycling has been confirmed by image analysis and used to help define proteins needed for post-endocytic recycling to the PM (MacDonald and Piper, 2017; Laidlaw *et al*., 2022a; Laidlaw *et al*., 2022b). In this study, we sought to further assess cis- and *trans*-acting factors in the endocytosis and recycling of Ste3 using live cell imaging with FAP-tagged Ste3.

We showed that FAP-tagged Ste3 was a functional receptor and could activate the mating pathway in *MATα* cells similar to WT Ste3 or Ste3-GFP (Figure 6). Past elegant biochemical approaches defined the Ste3 residues required for its efficient endocytosis post ligand addition (Roth and Davis, 1996; Roth *et al*., 1998; Chen and Davis, 2000; Roth and Davis, 2000; Chen and Davis, 2002). However, these experiments relied on cell fixation and cellular fractionation and thus did not monitor cell surface populations of Ste3 in real-time. The FAP imaging technique enabled us to map Ste3 trafficking dynamics with increased resolution to provide more detailed spatial information. We found that the bulk of Ste3 internalization occurs within the first 10 to 20 minutes following exposure to a-factor, rather than on the longer timeframes (40-120 min) reported previously (Chen and Davis, 2000). Interestingly, we noted that FAP-Ste3 was not evenly distributed at the PM but had a somewhat punctate distribution in the membrane. This is similar to the punctate patterning described for many PM proteins, which can partition into discrete PM subdomains with unique protein and lipid compositions (Lingwood and Simons, 2010; Schuberth and Wedlich-Soldner, 2015; Sezgin *et al*., 2017). It will be interesting in future studies to see which, if any, subdomain Ste3 may partition to at the cell surface.

Consistent with earlier findings, we demonstrated that the Ste3 C-terminal tail was essential for endocytosis (Roth and Davis, 1996; Roth *et al*., 1998; Roth and Davis, 2000). Within the tail, the PEST region was posited to be required only for steady-state receptor turnover (Roth *et al*., 1998; Roth and Davis, 2000). However, we find that disruption of sequences within the PEST motif similarly blocks ligand-induced endocytosis of Ste3. Three key lysines in the PEST motif are ubiquitinated to permit Ste3 endocytosis (Roth and Davis, 1996, 2000). We find that mutation of just one of these lysines, K424, is sufficient to disrupt ligand-induced Ste3 endocytosis, causing PM retention of a significant receptor population even after 60 min of ligand exposure. Past studies showed that Ste3 ubiquitination depends on Rsp5 (Rotin *et al*., 2000; Chen and Davis, 2002; Abazari *et al*., 2015; Prosser *et al*., 2015), yet Ste3 lacks Rsp5 interaction motifs. Indeed, we previously identified α-arrestins as important regulators of Rsp5-dependent Ste3 basal internalization via clathrin-mediated endocytosis (Prosser *et al*., 2015).

When comparing the trafficking of FAP- and pHluorin-tagged Ste3, we found that the Ste3 synthesized in response to a-factor accumulates at the shmoo tip about one hour after a-factor exposure (Figure 11F). Previous work detailing the transcriptional responses to a-factor did not elaborate on the localization of newly synthesized receptors (Hagen and Sprague, 1984). This increase in Ste3 abundance due to a-factor-induced transcriptional activation of Ste3 obscures endocytic dynamics when using fluorescent tags such as pHluorin that label the complete pool of Ste3 in cells. In contrast, the ability to selectively visualize distinct spatial and temporal pools of Ste3 using the FAP tag, in combination with the cell-impermeant dye, highlights the power of the imaging technology reported here. Using differentially tagged Ste3 populations, we showed that Ste3 present in the cell prior to a-factor addition and the newly synthesized Ste3 are targeted to spatially distinct PM locations (Figure 11F). Our data suggest a sorting mechanism that distinguishes “old” from “new” Ste3 and directs the receptor based on its past location in the cell.

What features dictate Ste3 post-endocytic sorting? Ste3 has been used as a model recycling cargo thanks to work that helped dissect factors needed for early endosomal recycling in yeast (MacDonald and Piper, 2017). These analyses were performed under basal, but not ligand-induced conditions, and identified Rcy1 (an F-box protein involved in endosomal recycling), Ist1 (a protein needed for endosomal recycling), and Nhx1 (a sodium/potassium exchanger needed for vacuole fusion) as endosomal factors that facilitate Ste3 recycling to the PM (MacDonald and Piper, 2017; Laidlaw *et al*., 2022a). Another study employing a deubiquitinase (DUb) fusion to fluorescently tagged Ste3 suggests that Gpa1, the Gα subunit of the mating pathway, and components of the glucose sensing machinery (e.g., the glucose sensing GPCR, Gpr1; a regulatory subunit of the Glc7 phosphatases, Reg1; and two transcriptional repressors, Mig1and Mig2) are also necessary for efficient recycling of Ste3 to the PM (Laidlaw *et al*., 2022b). In future studies, we will explore the role of these recycling factors with FAP-Ste3 and Ste3-pHluorin to evaluate their impact on Ste3 endocytosis and recycling in response to a-factor, thereby providing a real-time analysis of their contributions.

### α-Arrestins operate in both basal and ligand-induced GPCR endocytosis

In addition to defining the role of key *cis*-acting sequences in ligand-induced Ste3 trafficking that reside at the ‘tail’, we identify the α-arrestins required for PM turnover of Ste3. Consistent with our early studies of constitutive turnover of Ste3 (Prosser *et al*., 2015), we find that three α-arrestins, Aly1, Aly2, and Art1, are needed for ligand-induced Ste3 internalization. Here, we find that both constitutive and ligand-induced endocytosis of Ste3 is impaired in the absence of Art1, Aly1, and Aly2 (Figure 11F). However, these α-arrestins do not appear to impact Ste3 equivalently. In cells lacking Art1, Ste3 retention at the PM post a-factor treatment is robust and the receptor is evenly distributed across the cell surface (Figures 9 and S6). In contrast, in cells lacking Aly1/Aly2, Ste3 PM retention is less striking and shorter-lived (Figure 9). In addition, concentrated patches of Ste3 arise in *aly1*Δ *aly2*Δ cells, suggesting either that the Aly adaptors cannot internalize Ste3 from a subdomain of the PM or that Ste3 recycles to specific PM regions more effectively in the absence of these α-arrestins, resulting in patches of receptor close to bud neck or shmoo sites. These findings raise the possibility of functional partitioning for α-arrestins, which might allow them to regulate the trafficking of discrete pools of PM or intracellular receptors. More generally, numerous cargos are internalized by α-arrestins, yet mechanisms underlying the functional redundancy for these trafficking adaptors remain poorly understood.

The role of α-arrestins in regulating mammalian GPCRs remains somewhat controversial (Shea *et al*., 2012; Aubry and Klein, 2013; Han *et al*., 2013). However, their role in regulating GPCRs in yeast is established (Alvaro *et al*., 2014; Prosser *et al*., 2015; Emmerstorfer-Augustin *et al*., 2018). It would be surprising if the mammalian α-arrestins could not similarly regulate GPCRs, especially in light of the high-content studies that have identified many β-arrestin-independent GPCRs (Moo *et al*., 2021). Notably, mammalian α-arrestins have been shown to engage GPCRs, but for the two best-described examples to date, the α-arrestins are not operating in controlling endocytosis of these GPCRs. More specifically, mammalian α-arrestin ARRDC3 interacts with the β2-adrenergic receptor at endosomes in a ligand-independent manner to control the intracellular recycling of this receptor (Tian *et al*., 2016) but can also help drive ubiquitination of protease-activated receptor 1 (PAR1) to stimulate PAR1’s lysosomal degradation (Dores *et al*., 2015). These findings support the idea that human α-arrestins are likely able to engage GPCRs, though the precise sequences needed for α-arrestin-GPCR interaction have not been mapped to date.

In yeast, specific sequences in the Ste3 C-terminal tail are needed for its endocytosis. The defects in ligand-dependent endocytosis we observed for Ste3 C-tail mutants could be due to the elimination of post-translational regulatory sites or disruption of the binding sites for endocytic regulators, such as the α-arrestins and/or clathrin-binding adaptor proteins. The amino acids needed for α-arrestin-GPCR association have not yet been mapped. Considering that three α-arrestins mediate the internalization of Ste2 and Ste3 (*i.e.,* Rod1, Rog3, and Art1 or Aly1, Aly2, and Art1, respectively), there is an opportunity for both a common regulatory sequence (*i.e.,* for Art1) and divergent regulatory sequences (*i.e.,* for Rod1/Rog3 vs Aly1/Aly2) in Ste2 and Ste3. To date, only a few α-arrestin interaction interfaces have been defined for any membrane protein, but based on those, it appears that α-arrestins preferentially associate with acidic patches on cargo proteins (Lin *et al*., 2008; Wawrzycka *et al*., 2019; Ivashov *et al*., 2020; Barata-Antunes *et al*., 2022). Our earlier work demonstrated that α-arrestins can bind to the C-tails of Ste3 and Ste2, suggesting that key interactions are needed to recruit α-arrestins in these regions (Alvaro *et al*., 2014; Prosser *et al*., 2015). With FAP technology, we are well-positioned to define how α-arrestins recognize GPCRs, filling a significant knowledge gap in the field and further establishing α-arrestins as bona fide regulators of GPCR trafficking and signaling.

## MATERIALS AND METHODS

### Yeast strains and growth conditions

Yeast strains are listed in Supplemental Table 1 and derived from the BY4742 (*MATα*) genetic background of *S. cerevisiae* (S288C in origin). Deletion strains were built using a PCR-based method described previously (Longtine *et al*., 1998). Yeast cells were grown at 30°C and cultured in synthetic complete (SC) medium (2% glucose, yeast nitrogen base without amino acids, supplemented with amino acid drop-out mixtures for selection) or yeast extract peptone dextrose medium (YPD) (Johnston *et al*., 1977). Liquid medium was filter sterilized, and for plate medium, 2% w/v agar was added before autoclaving. Yeast cultures were grown overnight, then reinoculated (A_600_=0.2 or 0.3) and grown to mid-exponential log phase (∼4 hours to reach an A_600_=0.8*-*1.0) before experimentation.

### Plasmids and DNA manipulations

Plasmids used in this work are listed in Supplemental Table 2. Plasmid constructs were built using PCR amplification with Phusion High Fidelity DNA polymerase (ThermoFisher Scientific, Waltham, MA) and sequence validated through Sanger sequencing (Genewiz, South Plainfield, NJ). Plasmid maps were generated using SnapGene software (Insightful Science, Chicago, IL). Gene Blocks of the original and optimized sequence of FAP were obtained from GeneWiz (GeneWiz, South Plainfield, NJ). Codon optimization was done using the JAVA Codon Adaptation Tool (JCat) (Technical University of Braunschweig, Brunswick, Germany) (Grote *et al*., 2005). JCat employs a Codon Adaption Index (CAI) to generate an optimized DNA sequence by measuring the codon usage bias, which is the frequency of synonymous codon occurrence and cognate tRNAs usage in an organism (Grote *et al*., 2005). The FAP-tagging plasmids and those expressing these FAP-tagged cellular markers are all available on Addgene (see Supplemental Table 2 or Addgene global deposit number 84326). Plasmids were transformed into yeast cells using the lithium acetate method (Ausubel, 1991) and selected using SC media lacking specific amino acids or YPD medium supplemented with antibiotics.

### BSA coated tubes

The a-factor peptide is hydrophobic and tends to adhere to glass surfaces during purification. To ensure a-factor remained in solution, all tubes for a-factor treatments were coated with 1% w/v bovine serum albumin (BSA; Sigma-Aldrich, Waltham, MA) solution. Tubes were coated by filling them with the BSA solution and incubating at room temperature with rotation overnight. Following incubation, the BSA solution was removed, and tubes were rinsed with sterile water before use.

### Shmoo morphology assessment

To evaluate yeast morphology before and after a-factor treatment, cells were grown to saturation in SC medium at pH 6.6, inoculated in fresh medium (A_600_=0.3), and grown to mid-exponential log phase (∼A_600_=0.8). Equal densities of cells were collected by centrifugation and resuspended in 1 ml of fresh SC medium. Cells were plated onto concanavalin A-coated (MP Biomedicals, Solon OH, USA) 35 mm glass bottom microwell dishes (MatTek Corp., Ashland, MA) and the zero time-point was imaged using differential interference contrast (DIC) microscopy on a Nikon Ti inverted microscope (Nikon Instruments Ltd., Tokyo, Japan). Cells were transferred into BSA-coated glass culture tubes, 5 μM of a-factor mating pheromone (Zymo Research; 1mg/ml working stock) was added, and cells were incubated at 30°C for 4 hours prior to imaging and assessing shmoo formation.

The morphological composition of the cell populations was assessed using FIJI 2.0.0, a version of ImageJ (NIH, Bethesda, MA). Cells were assigned into “yeast-form” or “shmoo” populations using the cell counter tool. The percentage of cells with shmoo morphology was calculated, and statistical significance was determined using a Student’s t-test in Prism Software, version 10 (GraphPad, Boston, MA).

#### FAP Staining and Confocal Fluorescence Microscopy

To determine the localization and abundance of FAP-tagged Ste3, cells were grown as described above. Equal densities of cells (∼A_600_ 0.8) were collected by centrifugation and resuspended in 98 μl of fresh SC medium. A final concentration of 10 μM Cell Tracker Blue CMAC (7-amino-4-chloromethylcoumarin) dye (Life Technologies; Carlsbad, CA) and 1 μM of MG-TAU dye (cell permeant dye αRed-np1, Spectra Genetics, Pittsburgh, PA; 1:100 dilution of stock) or 1 μM of MG-ESTER (cell permeant dye αRed-p1, Spectra Genetics; 1:200 dilution of stock) was added to visualize vacuoles or the FAP tag at the PM or in whole cells, respectively. Cells with dye were incubated at 30°C with agitation for 15 minutes. Following incubation, cells were washed twice in 1 ml of fresh medium and transferred to a new BSA-coated tube. The zero timepoint was imaged from these cells, then 5 μM a-factor (Zymo Research, Orange, CA) was added to the rest of the cells, and they were incubated at 30°C with shaking, and sampled and imaged at the indicated timepoints. To image cells, 75 μl of cell suspension with 75 μl of fresh medium were plated onto 35 mm glass bottom microwell dishes (MatTek Corp., Ashland, MA) coated with concanavalin A (MP Biomedicals, Solon OH, USA). It is important to note that while malachite green itself can be toxic to yeast cells, the MG-derived dyes are not; even for cells incubated in these MG-derived dyes for prolonged periods (*i.e.,* ∼20 hours), there is no inhibition of yeast cell growth, unlike the robust inhibition of yeast cell growth that occurs when the same concentration of malachite green is used to treat cells (Szent-Gyorgyi *et al*., 2008).

For pHluorin imaging, cells were grown overnight to saturation in SC media, inoculated in fresh medium (A_600_=0.2) and grown to mid-exponential log phase (∼A_600_=0.8). Equal densities of cells were collected by centrifugation and resuspended in fresh SC medium, to which 10 μM CMAC dye was added to visualize vacuoles and cells were visualized post a-factor addition as indicated above. Cells were similarly incubated with MG-ESTER or MG-TAU dyes where simultaneous pHluorin and FAP-tagged Ste3 imaging was performed.

All confocal fluorescence microscopy was performed on a Nikon Eclipse Ti inverted microscope (Nikon Instruments Inc, Tokyo, Japan) outfitted with a swept field confocal scan head (Prairie Instruments, Middleton, WI), an Apo 100X objective (NA 1.49), an Agilent monolith laser launch (Agilent Technologies, Santa Clara, CA), and an Andor iXon3 camera (Oxford Instruments, Andor Technologies, Belfast, Northern Ireland). All images within an experiment were captured using identical parameters as controlled by the NIS-Elements software (Nikon Instruments Inc.). Images for figures were adjusted evenly within a figure panel, unless otherwise indicated, using NIS-Elements.

### Flow cytometry measurements

To quantitatively assess FAP fluorescence in dynamic populations, yeast cells containing the indicated FAP-tagged marker were cultured to mid-log phase (as indicated above) and fluorescence was analyzed using an Attune Nxt Flow Cytometer (ThermoFisher Scientific). Flow rate was set to collect 12.5 μl/min or 100,0000 events. The RL2 laser was used for excitation of the FAP probe and detected using the 720/730 filter. A threshold of 25x1000 was applied to the forward scatter to capture yeast cells. Data was analyzed using FlowJo software (Becton, Dickinson & Company, Franklin Lakes, NJ) to gate samples in forward and side scatter to include only single yeast cell events when measuring fluorescent values.

To define the pH sensitivity of the FAP-fluorescence, cells were grown in pH 6.6 medium overnight, inoculated into fresh pH 6.6 medium (A_600_=0.2) and grown for 2 h at 30°C. Cells were then collected by centrifugation, washed, and shifted to media of differing pH (adjusted to 4.1, 4.6, 5.1, 5.8, or 6.6 using hydrochloric acid or sodium hydroxide) and grown for 2 h in these new pH conditions. Equal densities of cells (∼A_600_=0.8) were collected by centrifugation and resuspended in 99 μl of fresh media of the corresponding pH for that experiment and 1 μM of either MG-TAU or MG-ESTER dye was added. Cells were incubated for 15 min at 30°C with agitation, washed twice, and resuspended in 1 ml of fresh media and placed in a new tube. Fluorescence was measured using an Attune NxT Flow Cytometer (ThermoFisher Scientific). The voltage settings for acquisition were 215 for FSC, 325 for SSC, and 450 for RL2. Cells that were not stained with the FAP dye were used to gate non-fluorescent cells.

### Quantitative imaging analyses and statistical tests

All images were manually quantified using FIJI software version 2.0.0 to measure fluorescence intensities. A 2-pixel wide line was hand-drawn around the perimeter of the plasma membrane (PM) to establish the ROI. The ROI was overlayed on the 405 channel to validate the vacuole co-staining (CMAC). Mean pixel intensity of the background was subtracted from each ROI as described (O’Donnell *et al*., 2015). At least three full imaging fields and 75 cells were analyzed for each dataset. Though hundreds to thousands of cells were analyzed for an individual experiment, at least three biological replicate experiments were performed for each condition. These data were collated into a Superplot (Prism Software, GraphPad) (Lord *et al*., 2020). The resulting data were evaluated using Prism software, where statistics were performed on the mean values from the three replicate experiments and not on the full population distributions. Kruskal-Wallis statistical analyses with Dunn’s post hoc test were used to define significant changes, denoted by *p value <0.1, **p value <0.01, ***p value <0.001, and ns p value >0.1.

To assess vacuolar fluorescence, the CMAC vacuolar stain was converted to a mask using FIJI software (as described in O’Donnell *et al*., 2015) that defined the vacuole regions of interest. This mask was then applied to the FAP channel (as for Figure 3G) and the median FAP signal in the vacuole measured. On occasion, peri-vacuolar compartments were included in this fluorescence measure as the CMAC mask was not always precise enough to differentiate between the vacuole and peri-vacuolar (likely MVBs) compartments, which may be docked onto the vacuole surface. Mean pixel intensity of the background was subtracted from each ROI as described (O’Donnell *et al*., 2015). Statistical analyses were performed using a Student’s t-test for Figure 3G.

To define the colocalization of FAP-tagged Ste3 with pHluorin-tagged Ste3 (Figures 11B and 11D), Pearson’s Correlation Coefficient was determined using the colocalizations plugin in FIJI software (Image J, NIH). To demonstrate the spatial distribution change between Ste3-pHluorin and FAP-Ste3, line scans were performed using the plot profile analysis tool in FIJI software (Image J, NIH) and plotting the values in Prism (GraphPad).

### Yeast protein extraction and immunoblot analysis

To analyze protein abundance, yeast whole-cell protein extracts were made by growing cells in SC medium with appropriate nutrient selection to mid-exponential phase at 30°C (A_600_ of 0.7-0.8) and then harvesting equivalent densities of cells by centrifugation. Cell pellets were flash-frozen in liquid nitrogen and stored at -80°C. To make extracts, pelleted cells were lysed, and proteins precipitated using the trichloroacetic acid extraction method as described (Volland *et al*., 1994). Protein precipitates were solubilized in SDS/urea sample buffer (40 mM Tris [pH 6.8], 0.1 mM EDTA, 5% SDS, 8 M urea, and 1% β-mercaptoethanol) (O’Donnell *et al*., 2013) and heated to 37°C for 15 min. Extracts were resolved by SDS-PAGE, and proteins were identified by immunoblotting. Either Revert^TM^ 700 Total Protein stain (LI-COR BioSciences, Lincoln, NE) of the membranes or anti-Zwf1 antibody (MilliporeSigma, St. Louis, MO) was used as a loading and transfer control. For immunoblotting, primary antibodies against Myc (cat # MA1980, Thermo Scientific, Waltham, MA), RFP (cat # 600-401-379, Rockland Immunochemicals, Inc., Pottstown, PA), ERK1/2 (C-9, sc 514302; Santa Cruz Biotechnology, Santa Cruz, CA), or Zwf1 (cat # A9521, MilliporeSigma, St. Louis, MO) were employed at the dilutions indicated from the manufacturers. Anti-mouse or anti-rabbit secondary antibodies conjugated to IRDye-800 or IRDye-680 (LI-COR BioSciences) were detected using the Odyssey CLx infrared imaging system (LI-COR BioSciences).

## ACKNOWLEDGMENTS

This research was funded by the National Sciences Foundation (MCB CAREER 1902859 and 1553143 to A.F.O., NSF 2321624 to A.F.O. and MCB CAREER 1942395 to D.C.P.) and the National Institutes of Health (R35 GM131732 to J.L.B., R01 HLB127711 grant to A.V.K.). The work was also supported by start-up funds from the Depts of Biological Sciences at Duquesne University and the University of Pittsburgh to A.F.O. The work was further supported by RRID SCR_022084 Microscopy Facility (C.S.) at the Univ. of Pittsburgh. The American Heart Association supported N.A.H. with a predoctoral fellowship. K.G.O. was supported by a Goldwater fellowship and the University of Pittsburgh’s Chancellor’s research fellowships. We thank Tova Finkelstein and Hillary Serbin for their work in generating the Ste3-GFP construct. We gratefully acknowledge the Brodsky and O’Donnell lab members for their helpful discussions and feedback on the work.

## AUTHOR CONTRIBUTIONS

K.G.O. performed most of the experiments presented in this work including Figures 2C-D, 3A-E and G, 6A-B, and 7-11 as well as Supplemental Figures S1, S2, S6-S7. K.G.O. also did much of the data analysis and assisted in writing and revising the manuscript. N.A.H. generated the plasmid constructs for FAP expression along with the FAP-tagged cellular markers, performed pilot experiments, and aided in initial versions of manuscript preparation. N.A.H. figure contributions include 1A-C, 2A-B and E-F, 3F, 4B-D, 5, and 6C-E. C.K.M. assisted in experiments, writing, and revising the manuscript. C.S. aided K.O. with the flow cytometry analyses and performed the quantification. E.F. built plasmids, performed sequence validation of constructs, generated plasmid map figures, and read and edited the manuscript. J.A.W. performed a subset of experiments with the optimized FAP tag. M.P.B. assisted with conceptual design and FAP technology implementation. J.L.B aided in conceptual design and edited the manuscript. D.C.P. generated the pHluorin constructs used and provided feedback on the conceptual design and manuscript editing. A.V.K. assisted with technical aspects for all the microscopy and quantification, and read and edited the manuscript. A.F.O. conceived of the project design, aided in experiment implementation, and wrote and revised the manuscript.

## Abbreviations

CFTR cystic fibrosis transmembrane conductance regulator

ER endoplasmic reticulum

FAP fluorogen activating protein

FP fluorescent proteins

GFP green fluorescent protein

GPCR G-protein coupled receptor

MAPK Mitogen activated protein kinase

MG malachite green

MG-Ester cell permeant MG-derived dye

MG-B-Tau cell impermeant MG-derived dye

MVB multi-vesicular body

PM plasma membrane

SCA single-chain antibody

WT wild-type

## REFERENCES

Abazari, A.M., Safavi, S.M., Rezazadeh, G., and Villanueva, L.G. (2015). Modelling the Size Effects on the Mechanical Properties of Micro/Nano Structures. Sensors (Basel) 15, 28543–28562.

Albert, B., Leger-Silvestre, I., Normand, C., Ostermaier, M.K., Perez-Fernandez, J., Panov, K.I., Zomerdijk, J.C., Schultz, P., and Gadal, O. (2011). RNA polymerase I-specific subunits promote polymerase clustering to enhance the rRNA gene transcription cycle. J Cell Biol 192, 277–293.

Alvarez, C.E. (2008). On the origins of arrestin and rhodopsin. BMC Evol Biol 8, 222.

Alvaro, C.G., O’Donnell, A.F., Prosser, D.C., Augustine, A.A., Goldman, A., Brodsky, J.L., Cyert, M.S., Wendland, B., and Thorner, J. (2014). Specific alpha-arrestins negatively regulate Saccharomyces cerevisiae pheromone response by down-modulating the G-protein-coupled receptor Ste2. Mol Cell Biol 34, 2660–2681.

Aridor, M., and Hannan, L.A. (2000). Traffic jam: a compendium of human diseases that affect intracellular transport processes. Traffic 1, 836–851.

Aubry, L., and Klein, G. (2013). True arrestins and arrestin-fold proteins: a structure-based appraisal. Prog Mol Biol Transl Sci 118, 21–56.

Ausubel, F.M. (1991). Current Protocols in Molecular Biology. John Wiley & Sons: New York.

Balaji, J., and Ryan, T.A. (2007). Single-vesicle imaging reveals that synaptic vesicle exocytosis and endocytosis are coupled by a single stochastic mode. Proc Natl Acad Sci U S A 104, 20576–20581.

Barata-Antunes, C., Talaia, G., Broutzakis, G., Ribas, D., De Beule, P., Casal, M., Stefan, C.J., Diallinas, G., and Paiva, S. (2022). Interactions of cytosolic tails in the Jen1 carboxylate transporter are critical for trafficking and transport activity. J Cell Sci 135.

Bardwell, L. (2005). A walk-through of the yeast mating pheromone response pathway. Peptides 26, 339–350.

Bardwell, L., Cook, J.G., Inouye, C.J., and Thorner, J. (1994). Signal propagation and regulation in the mating pheromone response pathway of the yeast Saccharomyces cerevisiae. Dev Biol 166, 363–379.

Becuwe, M., Vieira, N., Lara, D., Gomes-Rezende, J., Soares-Cunha, C., Casal, M., Haguenauer-Tsapis, R., Vincent, O., Paiva, S., and Leon, S. (2012). A molecular switch on an arrestin-like protein relays glucose signaling to transporter endocytosis. J Cell Biol 196, 247–259.

Boeck, J.M., and Spencer, J.V. (2017). Effect of human cytomegalovirus (HCMV) US27 on CXCR4 receptor internalization measured by fluorogen-activating protein (FAP) biosensors. PLoS One 12, e0172042.

Bohme, I., and Beck-Sickinger, A.G. (2009). Illuminating the life of GPCRs. Cell Commun Signal 7, 16.

Bonifacino, J.S., and Weissman, A.M. (1998). Ubiquitin and the control of protein fate in the secretory and endocytic pathways. Annu Rev Cell Dev Biol 14, 19–57.

Chen, I., Howarth, M., Lin, W., and Ting, A.Y. (2005). Site-specific labeling of cell surface proteins with biophysical probes using biotin ligase. Nat Methods 2, 99–104.

Chen, L., and Davis, N.G. (2000). Recycling of the yeast a-factor receptor. J Cell Biol 151, 731–738.

Chen, L., and Davis, N.G. (2002). Ubiquitin-independent entry into the yeast recycling pathway. Traffic 3, 110–123.

Chen, S., Webber, M.J., Vilardaga, J.P., Khatri, A., Brown, D., Ausiello, D.A., Lin, H.Y., and Bouley, R. (2011). Visualizing microtubule-dependent vasopressin type 2 receptor trafficking using a new high-affinity fluorescent vasopressin ligand. Endocrinology 152, 3893–3904.

Christianson, T.W., Sikorski, R.S., Dante, M., Shero, J.H., and Hieter, P. (1992). Multifunctional yeast high-copy-number shuttle vectors. Gene 110, 119–122.

Cole, S.R., Ashman, L.K., and Ey, P.L. (1987). Biotinylation: an alternative to radioiodination for the identification of cell surface antigens in immunoprecipitates. Mol Immunol 24, 699–705.

Coppinger, J.A., Hutt, D.M., Razvi, A., Koulov, A.V., Pankow, S., Yates, J.R., 3rd, and Balch, W.E. (2012). A chaperone trap contributes to the onset of cystic fibrosis. PLoS One 7, e37682.

Couve, A., and Hirsch, J.P. (1996). Loss of sustained Fus3p kinase activity and the G1 arrest response in cells expressing an inappropriate pheromone receptor. Mol Cell Biol 16, 4478–4485.

Deshaies, R.J., and Schekman, R. (1987). A yeast mutant defective at an early stage in import of secretory protein precursors into the endoplasmic reticulum. J Cell Biol 105, 633–645.

Dores, M.R., Lin, H., N, J.G., Mendez, F., and Trejo, J. (2015). The alpha-arrestin ARRDC3 mediates ALIX ubiquitination and G protein-coupled receptor lysosomal sorting. Mol Biol Cell 26, 4660–4673.

Dundas, C.M., Demonte, D., and Park, S. (2013). Streptavidin-biotin technology: improvements and innovations in chemical and biological applications. Appl Microbiol Biotechnol 97, 9343–9353.

Dunn, R., and Hicke, L. (2001). Multiple roles for Rsp5p-dependent ubiquitination at the internalization step of endocytosis. J Biol Chem 276, 25974–25981.

Emmerstorfer-Augustin, A., Augustin, C.M., Shams, S., and Thorner, J. (2018). Tracking yeast pheromone receptor Ste2 endocytosis using fluorogen-activating protein tagging. Mol Biol Cell 29, 2720–2736.

Feldhaus, M.J., Siegel, R.W., Opresko, L.K., Coleman, J.R., Feldhaus, J.M., Yeung, Y.A., Cochran, J.R., Heinzelman, P., Colby, D., Swers, J., Graff, C., Wiley, H.S., and Wittrup, K.D. (2003). Flow-cytometric isolation of human antibodies from a nonimmune Saccharomyces cerevisiae surface display library. Nat Biotechnol 21, 163–170.

Fernandez-Suarez, M., Baruah, H., Martinez-Hernandez, L., Xie, K.T., Baskin, J.M., Bertozzi, C.R., and Ting, A.Y. (2007). Redirecting lipoic acid ligase for cell surface protein labeling with small-molecule probes. Nat Biotechnol 25, 1483–1487.

Fisher, G.W., Adler, S.A., Fuhrman, M.H., Waggoner, A.S., Bruchez, M.P., and Jarvik, J.W. (2010). Detection and quantification of beta2AR internalization in living cells using FAP-based biosensor technology. J Biomol Screen 15, 703–709.

Gallo, E. (2020). Fluorogen-Activating Proteins: Next-Generation Fluorescence Probes for Biological Research. Bioconjug Chem 31, 16–27.

Gautier, A., Juillerat, A., Heinis, C., Correa, I.R., Jr., Kindermann, M., Beaufils, F., and Johnsson, K. (2008). An engineered protein tag for multiprotein labeling in living cells. Chem Biol 15, 128–136.

Glick, B.S., and Nakano, A. (2009). Membrane traffic within the Golgi apparatus. Annu Rev Cell Dev Biol 25, 113–132.

Goeckeler-Fried, J.L., Aldrin Denny, R., Joshi, D., Hill, C., Larsen, M.B., Chiang, A.N., Frizzell, R.A., Wipf, P., Sorscher, E.J., and Brodsky, J.L. (2021). Improved correction of F508del-CFTR biogenesis with a folding facilitator and an inhibitor of protein ubiquitination. Bioorg Med Chem Lett 48, 128243.

Goh, L.K., Huang, F., Kim, W., Gygi, S., and Sorkin, A. (2010). Multiple mechanisms collectively regulate clathrin-mediated endocytosis of the epidermal growth factor receptor. J Cell Biol 189, 871–883.

Griffin, B.A., Adams, S.R., and Tsien, R.Y. (1998). Specific covalent labeling of recombinant protein molecules inside live cells. Science 281, 269–272.

Grote, A., Hiller, K., Scheer, M., Munch, R., Nortemann, B., Hempel, D.C., and Jahn, D. (2005). JCat: a novel tool to adapt codon usage of a target gene to its potential expression host. Nucleic Acids Res 33, W526–531.

Hagen, D.C., and Sprague, G.F., Jr. (1984). Induction of the yeast alpha-specific STE3 gene by the peptide pheromone a-factor. J Mol Biol 178, 835–852.

Hager, N.A., Krasowski, C.J., Mackie, T.D., Kolb, A.R., Needham, P.G., Augustine, A.A., Dempsey, A., Szent-Gyorgyi, C., Bruchez, M.P., Bain, D.J., Kwiatkowski, A.V., O’Donnell, A.F., and Brodsky, J.L. (2018). Select alpha-arrestins control cell-surface abundance of the mammalian Kir2.1 potassium channel in a yeast model. J Biol Chem 293, 11006–11021.

Hager, N.A., McAtee, C.K., Lesko, M.A., and O’Donnell, A.F. (2021). Inwardly Rectifying Potassium Channel Kir2.1 and its “Kir-ious” Regulation by Protein Trafficking and Roles in Development and Disease. Front Cell Dev Biol 9, 796136.

Han, S.O., Kommaddi, R.P., and Shenoy, S.K. (2013). Distinct roles for beta-arrestin2 and arrestin-domain-containing proteins in beta2 adrenergic receptor trafficking. EMBO Rep 14, 164–171.

Herrador, A., Leon, S., Haguenauer-Tsapis, R., and Vincent, O. (2013). A mechanism for protein monoubiquitination dependent on a trans-acting ubiquitin-binding domain. J Biol Chem 288, 16206–16211.

Hicke, L., Zanolari, B., and Riezman, H. (1998). Cytoplasmic tail phosphorylation of the alpha-factor receptor is required for its ubiquitination and internalization. J Cell Biol 141, 349–358.

Holleran, J., Brown, D., Fuhrman, M.H., Adler, S.A., Fisher, G.W., and Jarvik, J.W. (2010). Fluorogen-activating proteins as biosensors of cell-surface proteins in living cells. Cytometry A 77, 776–782.

Holleran, J.P., Glover, M.L., Peters, K.W., Bertrand, C.A., Watkins, S.C., Jarvik, J.W., and Frizzell, R.A. (2012). Pharmacological rescue of the mutant cystic fibrosis transmembrane conductance regulator (CFTR) detected by use of a novel fluorescence platform. Mol Med 18, 685–696.

Howell, G.J., Holloway, Z.G., Cobbold, C., Monaco, A.P., and Ponnambalam, S. (2006). Cell biology of membrane trafficking in human disease. Int Rev Cytol 252, 1–69.

Huang, F., Goh, L.K., and Sorkin, A. (2007). EGF receptor ubiquitination is not necessary for its internalization. Proc Natl Acad Sci U S A 104, 16904–16909.

Huh, W.K., Falvo, J.V., Gerke, L.C., Carroll, A.S., Howson, R.W., Weissman, J.S., and O’Shea, E.K. (2003). Global analysis of protein localization in budding yeast. Nature 425, 686–691.

Hung, M.C., and Link, W. (2011). Protein localization in disease and therapy. J Cell Sci 124, 3381–3392.

Ivashov, V., Zimmer, J., Schwabl, S., Kahlhofer, J., Weys, S., Gstir, R., Jakschitz, T., Kremser, L., Bonn, G.K., Lindner, H., Huber, L.A., Leon, S., Schmidt, O., and Teis, D. (2020). Complementary alpha-arrestin-ubiquitin ligase complexes control nutrient transporter endocytosis in response to amino acids. Elife 9.

Johnston, G.C., Pringle, J.R., and Hartwell, L.H. (1977). Coordination of growth with cell division in the yeast Saccharomyces cerevisiae. Exp Cell Res 105, 79–98.

Jonker, C.T.H., Deo, C., Zager, P.J., Tkachuk, A.N., Weinstein, A.M., Rodriguez-Boulan, E., Lavis, L.D., and Schreiner, R. (2020). Accurate measurement of fast endocytic recycling kinetics in real time. J Cell Sci 133.

Kane, P.M. (1999). Biosynthesis and regulation of the yeast vacuolar H+-ATPase. J Bioenerg Biomembr 31, 49–56.

Kane, P.M. (2006). The where, when, and how of organelle acidification by the yeast vacuolar H+-ATPase. Microbiol Mol Biol Rev 70, 177–191.

Keppler, A., Gendreizig, S., Gronemeyer, T., Pick, H., Vogel, H., and Johnsson, K. (2003). A general method for the covalent labeling of fusion proteins with small molecules in vivo. Nat Biotechnol 21, 86–89.

Keppler, A., Pick, H., Arrivoli, C., Vogel, H., and Johnsson, K. (2004). Labeling of fusion proteins with synthetic fluorophores in live cells. Proc Natl Acad Sci U S A 101, 9955–9959.

Kerjan, P., Cherest, H., and Surdin-Kerjan, Y. (1986). Nucleotide sequence of the Saccharomyces cerevisiae MET25 gene. Nucleic Acids Res 14, 7861–7871.

Klis, F.M., Mol, P., Hellingwerf, K., and Brul, S. (2002). Dynamics of cell wall structure in Saccharomyces cerevisiae. FEMS Microbiol Rev 26, 239–256.

Komatsu, T., Johnsson, K., Okuno, H., Bito, H., Inoue, T., Nagano, T., and Urano, Y. (2011). Real-time measurements of protein dynamics using fluorescence activation-coupled protein labeling method. J Am Chem Soc 133, 6745–6751.

Koulov, A.V., LaPointe, P., Lu, B., Razvi, A., Coppinger, J., Dong, M.Q., Matteson, J., Laister, R., Arrowsmith, C., Yates, J.R., 3rd, and Balch, W.E. (2010). Biological and structural basis for Aha1 regulation of Hsp90 ATPase activity in maintaining proteostasis in the human disease cystic fibrosis. Mol Biol Cell 21, 871–884.

Krysan, D.J., Ting, E.L., Abeijon, C., Kroos, L., and Fuller, R.S. (2005). Yapsins are a family of aspartyl proteases required for cell wall integrity in Saccharomyces cerevisiae. Eukaryot Cell 4, 1364–1374.

Labbe, S., and Thiele, D.J. (1999). Copper ion inducible and repressible promoter systems in yeast. Methods Enzymol 306, 145–153.

Laidlaw, K.M.E., Calder, G., and MacDonald, C. (2022a). Recycling of cell surface membrane proteins from yeast endosomes is regulated by ubiquitinated Ist1. J Cell Biol 221.

Laidlaw, K.M.E., Paine, K.M., Bisinski, D.D., Calder, G., Hogg, K., Ahmed, S., James, S., O’Toole, P.J., and MacDonald, C. (2022b). Endosomal cargo recycling mediated by Gpa1 and phosphatidylinositol 3-kinase is inhibited by glucose starvation. Mol Biol Cell 33, ar31.

Lee, L.G., Chen, C.H., and Chiu, L.A. (1986). Thiazole orange: a new dye for reticulocyte analysis. Cytometry 7, 508–517.

Leng, S., Qiao, Q., Miao, L., Deng, W., Cui, J., and Xu, Z. (2017). A wash-free SNAP-tag fluorogenic probe based on the additive effects of quencher release and environmental sensitivity. Chem Commun (Camb) 53, 6448–6451.

Li, Y., Kane, T., Tipper, C., Spatrick, P., and Jenness, D.D. (1999). Yeast mutants affecting possible quality control of plasma membrane proteins. Mol Cell Biol 19, 3588–3599.

Lin, C.H., MacGurn, J.A., Chu, T., Stefan, C.J., and Emr, S.D. (2008). Arrestin-related ubiquitin-ligase adaptors regulate endocytosis and protein turnover at the cell surface. Cell 135, 714–725.

Lingwood, D., and Simons, K. (2010). Lipid rafts as a membrane-organizing principle. Science 327, 46–50.

Lippincott-Schwartz, J., Altan-Bonnet, N., and Patterson, G.H. (2003). Photobleaching and photoactivation: following protein dynamics in living cells. Nat Cell Biol Suppl, S7–14.

Longtine, M.S., McKenzie, A., 3rd, Demarini, D.J., Shah, N.G., Wach, A., Brachat, A., Philippsen, P., and Pringle, J.R. (1998). Additional modules for versatile and economical PCR-based gene deletion and modification in Saccharomyces cerevisiae. Yeast 14, 953–961.

Lopes-Pacheco, M. (2016). CFTR Modulators: Shedding Light on Precision Medicine for Cystic Fibrosis. Front Pharmacol 7, 275.

Lopes-Pacheco, M. (2019). CFTR Modulators: The Changing Face of Cystic Fibrosis in the Era of Precision Medicine. Front Pharmacol 10, 1662.

Lord, S.J., Velle, K.B., Mullins, R.D., and Fritz-Laylin, L.K. (2020). SuperPlots: Communicating reproducibility and variability in cell biology. J Cell Biol 219.

Lorenz-Guertin, J.M., Wilcox, M.R., Zhang, M., Larsen, M.B., Pilli, J., Schmidt, B.F., Bruchez, M.P., Johnson, J.W., Waggoner, A.S., Watkins, S.C., and Jacob, T.C. (2017). A versatile optical tool for studying synaptic GABAA receptor trafficking. J Cell Sci 130, 3933–3945.

Los, G.V., Encell, L.P., McDougall, M.G., Hartzell, D.D., Karassina, N., Zimprich, C., Wood, M.G., Learish, R., Ohana, R.F., Urh, M., Simpson, D., Mendez, J., Zimmerman, K., Otto, P., Vidugiris, G., Zhu, J., Darzins, A., Klaubert, D.H., Bulleit, R.F., and Wood, K.V. (2008). HaloTag: a novel protein labeling technology for cell imaging and protein analysis. ACS Chem Biol 3, 373–382.

Luedtke, N.W., Dexter, R.J., Fried, D.B., and Schepartz, A. (2007). Surveying polypeptide and protein domain conformation and association with FlAsH and ReAsH. Nat Chem Biol 3, 779–784.

MacDonald, C., and Piper, R.C. (2017). Genetic dissection of early endosomal recycling highlights a TORC1-independent role for Rag GTPases. J Cell Biol 216, 3275–3290.

MacDonald, C., Shields, S.B., Williams, C.A., Winistorfer, S., and Piper, R.C. (2020). A Cycle of Ubiquitination Regulates Adaptor Function of the Nedd4-Family Ubiquitin Ligase Rsp5. Curr Biol 30, 465–479 e465.

Masuoka, J., Guthrie, L., and Hazen, K. (2002). Complications in cell-surface labelling by biotinylation of Candida albicans due to avidin conjugate binding to cell-wall proteins. Microbiology, 1073–1079.

Meurer, M., Duan, Y., Sass, E., Kats, I., Herbst, K., Buchmuller, B.C., Dederer, V., Huber, F., Kirrmaier, D., Stefl, M., Van Laer, K., Dick, T.P., Lemberg, M.K., Khmelinskii, A., Levy, E.D., and Knop, M. (2018). Genome-wide C-SWAT library for high-throughput yeast genome tagging. Nat Methods 15, 598–600.

Miyawaki, A., Sawano, A., and Kogure, T. (2003). Lighting up cells: labelling proteins with fluorophores. Nat Cell Biol Suppl, S1–7.

Moo, E.V., van Senten, J.R., Brauner-Osborne, H., and Moller, T.C. (2021). Arrestin-Dependent and -Independent Internalization of G Protein-Coupled Receptors: Methods, Mechanisms, and Implications on Cell Signaling. Mol Pharmacol 99, 242–255.

Moskowitz, S.M., Chmiel, J.F., Sternen, D.L., Cheng, E., Gibson, R.L., Marshall, S.G., and Cutting, G.R. (2008). Clinical practice and genetic counseling for cystic fibrosis and CFTR-related disorders. Genet Med 10, 851–868.

Motizuki, M., Yokota, S., and Tsurugi, K. (2008). Effect of low pH on organization of the actin cytoskeleton in Saccharomyces cerevisiae. Biochim Biophys Acta 1780, 179–184.

Mukherjee, M., Nandi, A., Chandra, K., Saikia, S.K., Jana, C.K., and Das, N. (2020). Protein extraction from Saccharomyces cerevisiae at different growth phases. J Microbiol Methods 172, 105906.

Mumberg, D., Muller, R., and Funk, M. (1995). Yeast vectors for the controlled expression of heterologous proteins in different genetic backgrounds. Gene 156, 119–122.

Naganbabu, M., Perkins, L.A., Wang, Y., Kurish, J., Schmidt, B.F., and Bruchez, M.P. (2016). Multiexcitation Fluorogenic Labeling of Surface, Intracellular, and Total Protein Pools in Living Cells. Bioconjug Chem 27, 1525–1531.

Nicholson-Fish, J.C., Smillie, K.J., and Cousin, M.A. (2016). Monitoring activity-dependent bulk endocytosis with the genetically-encoded reporter VAMP4-pHluorin. J Neurosci Methods 266, 1–10.

Nikko, E., and Pelham, H.R. (2009). Arrestin-mediated endocytosis of yeast plasma membrane transporters. Traffic 10, 1856–1867.

Nishimura, N., and Sasaki, T. (2008). Cell-surface biotinylation to study endocytosis and recycling of occludin. Methods Mol Biol 440, 89–96.

O’Donnell, A.F., Apffel, A., Gardner, R.G., and Cyert, M.S. (2010). Alpha-arrestins Aly1 and Aly2 regulate intracellular trafficking in response to nutrient signaling. Mol Biol Cell 21, 3552–3566.

O’Donnell, A.F., Huang, L., Thorner, J., and Cyert, M.S. (2013). A calcineurin-dependent switch controls the trafficking function of alpha-arrestin Aly1/Art6. J Biol Chem 288, 24063–24080.

O’Donnell, A.F., McCartney, R.R., Chandrashekarappa, D.G., Zhang, B.B., Thorner, J., and Schmidt, M.C. (2015). 2-Deoxyglucose impairs Saccharomyces cerevisiae growth by stimulating Snf1-regulated and alpha-arrestin-mediated trafficking of hexose transporters 1 and 3. Mol Cell Biol 35, 939–955.

O’Donnell, A.F., and Schmidt, M.C. (2019). AMPK-Mediated Regulation of Alpha-Arrestins and Protein Trafficking. Int J Mol Sci 20.

Perkins, L.A., and Bruchez, M.P. (2020). Fluorogen activating protein toolset for protein trafficking measurements. Traffic 21, 333–348.

Powell, C.D., Quain, D.E., and Smart, K.A. (2003). The impact of brewing yeast cell age on fermentation performance, attenuation and flocculation. FEMS Yeast Res 3, 149–157.

Pratt, C.P., He, J., Wang, Y., Barth, A.L., and Bruchez, M.P. (2015). Fluorogenic Green-Inside Red-Outside (GIRO) Labeling Approach Reveals Adenylyl Cyclase-Dependent Control of BKalpha Surface Expression. Bioconjug Chem 26, 1963–1971.

Prosser, D.C., Pannunzio, A.E., Brodsky, J.L., Thorner, J., Wendland, B., and O’Donnell, A.F. (2015). alpha-Arrestins participate in cargo selection for both clathrin-independent and clathrin-mediated endocytosis. J Cell Sci 128, 4220–4234.

Prosser, D.C., Whitworth, K., and Wendland, B. (2010). Quantitative analysis of endocytosis with cytoplasmic pHluorin chimeras. Traffic 11, 1141–1150.

Prosser, D.C., Wrasman, K., Woodard, T.K., O’Donnell, A.F., and Wendland, B. (2016). Applications of pHluorin for Quantitative, Kinetic and High-throughput Analysis of Endocytosis in Budding Yeast. J Vis Exp.

Roth, A.F., and Davis, N.G. (1996). Ubiquitination of the yeast a-factor receptor. J Cell Biol 134, 661–674.

Roth, A.F., and Davis, N.G. (2000). Ubiquitination of the PEST-like endocytosis signal of the yeast a-factor receptor. J Biol Chem 275, 8143–8153.

Roth, A.F., Sullivan, D.M., and Davis, N.G. (1998). A large PEST-like sequence directs the ubiquitination, endocytosis, and vacuolar degradation of the yeast a-factor receptor. J Cell Biol 142, 949–961.

Rotin, D., Staub, O., and Haguenauer-Tsapis, R. (2000). Ubiquitination and endocytosis of plasma membrane proteins: role of Nedd4/Rsp5p family of ubiquitin-protein ligases. J Membr Biol 176, 1–17.

Schandel, K.A., and Jenness, D.D. (1994). Direct evidence for ligand-induced internalization of the yeast alpha-factor pheromone receptor. Mol Cell Biol 14, 7245–7255.

Schuberth, C., and Wedlich-Soldner, R. (2015). Building a patchwork - The yeast plasma membrane as model to study lateral domain formation. Biochim Biophys Acta 1853, 767–774.

Sezgin, E., Levental, I., Mayor, S., and Eggeling, C. (2017). The mystery of membrane organization: composition, regulation and roles of lipid rafts. Nat Rev Mol Cell Biol 18, 361–374.

Shank, N.I., Pham, H.H., Waggoner, A.S., and Armitage, B.A. (2013). Twisted cyanines: a non-planar fluorogenic dye with superior photostability and its use in a protein-based fluoromodule. J Am Chem Soc 135, 242–251.

Shea, F.F., Rowell, J.L., Li, Y., Chang, T.H., and Alvarez, C.E. (2012). Mammalian alpha arrestins link activated seven transmembrane receptors to Nedd4 family e3 ubiquitin ligases and interact with beta arrestins. PLoS One 7, e50557.

Shih, S.C., Katzmann, D.J., Schnell, J.D., Sutanto, M., Emr, S.D., and Hicke, L. (2002). Epsins and Vps27p/Hrs contain ubiquitin-binding domains that function in receptor endocytosis. Nat Cell Biol 4, 389–393.

Shiwarski, D.J., Darr, M., Telmer, C.A., Bruchez, M.P., and Puthenveedu, M.A. (2017). PI3K class II alpha regulates delta-opioid receptor export from the trans-Golgi network. Mol Biol Cell 28, 2202–2219.

Sikorski, R.S., and Hieter, P. (1989). A system of shuttle vectors and yeast host strains designed for efficient manipulation of DNA in Saccharomyces cerevisiae. Genetics 122, 19–27.

Silva, G.L., Ediz, V., Yaron, D., and Armitage, B.A. (2007). Experimental and computational investigation of unsymmetrical cyanine dyes: understanding torsionally responsive fluorogenic dyes. J Am Chem Soc 129, 5710–5718.

Smith, A.E., Zhang, Z., Thomas, C.R., Moxham, K.E., and Middelberg, A.P. (2000). The mechanical properties of Saccharomyces cerevisiae. Proc Natl Acad Sci U S A 97, 9871–9874.

Sorkin, A., and Von Zastrow, M. (2002). Signal transduction and endocytosis: close encounters of many kinds. Nat Rev Mol Cell Biol 3, 600–614.

Sprague, G.F., Jr., Jensen, R., and Herskowitz, I. (1983). Control of yeast cell type by the mating type locus: positive regulation of the alpha-specific STE3 gene by the MAT alpha 1 product. Cell 32, 409–415.

Szent-Gyorgyi, C., Schmidt, B.F., Creeger, Y., Fisher, G.W., Zakel, K.L., Adler, S., Fitzpatrick, J.A., Woolford, C.A., Yan, Q., Vasilev, K.V., Berget, P.B., Bruchez, M.P., Jarvik, J.W., and Waggoner, A. (2008). Fluorogen-activating single-chain antibodies for imaging cell surface proteins. Nat Biotechnol 26, 235–240.

Szent-Gyorgyi, C., Stanfield, R.L., Andreko, S., Dempsey, A., Ahmed, M., Capek, S., Waggoner, A.S., Wilson, I.A., and Bruchez, M.P. (2013). Malachite green mediates homodimerization of antibody VL domains to form a fluorescent ternary complex with singular symmetric interfaces. J Mol Biol 425, 4595–4613.

Tanaka, T., Zhou, Y., Ozawa, T., Okizono, R., Banba, A., Yamamura, T., Oga, E., Muraguchi, A., and Sakurai, H. (2018). Ligand-activated epidermal growth factor receptor (EGFR) signaling governs endocytic trafficking of unliganded receptor monomers by non-canonical phosphorylation. J Biol Chem 293, 2288–2301.

Tham, D.K.L., and Moukhles, H. (2017). Determining Cell-surface Expression and Endocytic Rate of Proteins in Primary Astrocyte Cultures Using Biotinylation. J Vis Exp.

Tian, X., Irannejad, R., Bowman, S.L., Du, Y., Puthenveedu, M.A., von Zastrow, M., and Benovic, J.L. (2016). The alpha-Arrestin ARRDC3 Regulates the Endosomal Residence Time and Intracellular Signaling of the beta2-Adrenergic Receptor. J Biol Chem 291, 14510–14525.

Todorow, Z., Spang, A., Carmack, E., Yates, J., and Schekman, R. (2000). Active recycling of yeast Golgi mannosyltransferase complexes through the endoplasmic reticulum. Proc Natl Acad Sci U S A 97, 13643–13648.

Tomas, A., Futter, C.E., and Eden, E.R. (2014). EGF receptor trafficking: consequences for signaling and cancer. Trends Cell Biol 24, 26–34.

Toshima, J.Y., Nakanishi, J., Mizuno, K., Toshima, J., and Drubin, D.G. (2009). Requirements for recruitment of a G protein-coupled receptor to clathrin-coated pits in budding yeast. Mol Biol Cell 20, 5039–5050.

Toshima, J.Y., Nishinoaki, S., Sato, Y., Yamamoto, W., Furukawa, D., Siekhaus, D.E., Sawaguchi, A., and Toshima, J. (2014). Bifurcation of the endocytic pathway into Rab5-dependent and -independent transport to the vacuole. Nat Commun 5, 3498.

Toshima, J.Y., Toshima, J., Kaksonen, M., Martin, A.C., King, D.S., and Drubin, D.G. (2006). Spatial dynamics of receptor-mediated endocytic trafficking in budding yeast revealed by using fluorescent alpha-factor derivatives. Proc Natl Acad Sci U S A 103, 5793–5798.

Umezawa, H., Aoyagi, T., Morishima, H., Matsuzaki, M., and Hamada, M. (1970). Pepstatin, a new pepsin inhibitor produced by Actinomycetes. J Antibiot (Tokyo) 23, 259–262.

Urbanowski, J.L., and Piper, R.C. (2001). Ubiquitin sorts proteins into the intralumenal degradative compartment of the late-endosome/vacuole. Traffic 2, 622–630.

Valli, M., Sauer, M., Branduardi, P., Borth, N., Porro, D., and Mattanovich, D. (2005). Intracellular pH distribution in Saccharomyces cerevisiae cell populations, analyzed by flow cytometry. Appl Environ Microbiol 71, 1515–1521.

Volland, C., Galan, J.M., Urban-Grimal, D., Devilliers, G., and Haguenauer-Tsapis, R. (1994). Endocytose and degradation of the uracil permease of S. cerevisiae under stress conditions: possible role of ubiquitin. Folia Microbiol (Praha) 39, 554–557.

Watanabe, S., Mizukami, S., Hori, Y., and Kikuchi, K. (2010). Multicolor protein labeling in living cells using mutant beta-lactamase-tag technology. Bioconjug Chem 21, 2320–2326.

Wawrzycka, D., Sadlak, J., Maciaszczyk-Dziubinska, E., and Wysocki, R. (2019). Rsp5-dependent endocytosis and degradation of the arsenite transporter Acr3 requires its N-terminal acidic tail as an endocytic sorting signal and arrestin-related ubiquitin-ligase adaptors. Biochim Biophys Acta Biomembr 1861, 916–925.

Woolford, C.A., Daniels, L.B., Park, F.J., Jones, E.W., Van Arsdell, J.N., and Innis, M.A. (1986). The PEP4 gene encodes an aspartyl protease implicated in the posttranslational regulation of Saccharomyces cerevisiae vacuolar hydrolases. Mol Cell Biol 6, 2500–2510.

Yan, Q., Schmidt, B.F., Perkins, L.A., Naganbabu, M., Saurabh, S., Andreko, S.K., and Bruchez, M.P. (2015). Near-instant surface-selective fluorogenic protein quantification using sulfonated triarylmethane dyes and fluorogen activating proteins. Org Biomol Chem 13, 2078–2086.

Yao, J.Z., Uttamapinant, C., Poloukhtine, A., Baskin, J.M., Codelli, J.A., Sletten, E.M., Bertozzi, C.R., Popik, V.V., and Ting, A.Y. (2012). Fluorophore targeting to cellular proteins via enzyme-mediated azide ligation and strain-promoted cycloaddition. J Am Chem Soc 134, 3720–3728.

Yarwood, R., Hellicar, J., Woodman, P.G., and Lowe, M. (2020). Membrane trafficking in health and disease. Dis Model Mech 13.

Young, B.P., Craven, R.A., Reid, P.J., Willer, M., and Stirling, C.J. (2001). Sec63p and Kar2p are required for the translocation of SRP-dependent precursors into the yeast endoplasmic reticulum in vivo. EMBO J 20, 262–271.

Young, M.R., Heit, S., and Bublitz, M. (2024). Structure, function and biogenesis of the fungal proton pump Pma1. Biochim Biophys Acta Mol Cell Res 1871, 119600.

Zanolari, B., Raths, S., Singer-Kruger, B., and Riezman, H. (1992). Yeast pheromone receptor endocytosis and hyperphosphorylation are independent of G protein-mediated signal transduction. Cell 71, 755–763.

Zanolari, B., and Riezman, H. (1991). Quantitation of alpha-factor internalization and response during the Saccharomyces cerevisiae cell cycle. Mol Cell Biol 11, 5251–5258.

Zurn, A., Klenk, C., Zabel, U., Reiner, S., Lohse, M.J., and Hoffmann, C. (2010). Site-specific, orthogonal labeling of proteins in intact cells with two small biarsenical fluorophores. Bioconjug Chem 21, 853–859.

