## Supplemental for "Optimization of the fluorogen-activating protein tag for quantitative protein trafficking and co-localization studies in *S. cerevisiae*"

### Supporting Information

#### A new series of fluorogen-activating proteins for quantitative protein trafficking and co-localization studies in *S. cerevisiae*

Katherine G. Oppenheimer, Natalie A. Hager, Ceara K. McAtee, Elif Filiztekin, Chaowei Shang, Justina A. Warnick, Marcel P. Bruchez, Jeffrey L. Brodsky, Derek C. Prosser, Adam V. Kwiatkowski and Allyson F. O'Donnell.

**Supplemental Figure S1** (accompanies Figure 2). Negative gating for flow cytometry.

**Supplemental Figure S2** (accompanies Figure 3). FAP fluorescence response to pH changes.

**Supplemental Figure S3** (accompanies Figure 4). Inducing FAP expression from *MET25* or *CUP1* promoters.

**Supplemental Figure S4** (accompanies Figure 4). FAP-expression and -tagging plasmids.

**Supplemental Figure S5** (accompanies Figure 9). Contribution of $\alpha$-arrestins to steady-state turnover of FAP-Ste3.

**Supplemental Figure S6** (accompanies Figure 10). Contribution of $\alpha$-arrestins to steady-state turnover of Ste3-pHluorin.

**Supplemental Figure S7** (accompanies Figure 11). FAP-Ste3 can be used to monitor recycling from the PM.

**Supplemental Table 1.** Yeast strains used in this study.

**Supplemental Table 2.** Plasmids used in this study.

### Supplemental Figures & Legends

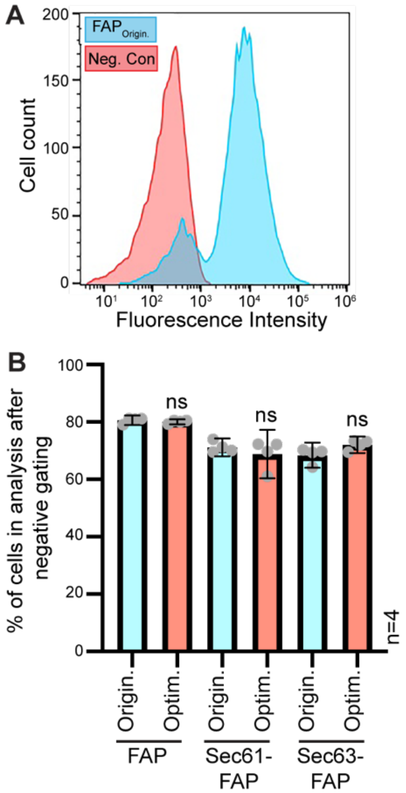

**Supplemental Figure S1 (accompanies Figure 2): Negative gating for flow cytometry.** (A) Cytometry analysis of WT cells either expressing soluble $FAP_{ORIGIN}$ or expressing no fluorophore (negative control for non-fluorescent cells). These non-fluorescent cells were used as a control to remove background cell autofluorescence or non-expressing cells (as occurs sometimes in small subpopulations when plasmids are used) by gating. (B) The percentage of cells remaining in each experiment performed for Figure 2D after the non-fluorescent cells were removed by gating, the threshold for which was established in panel S1A. In all cases, ~80% of cells remained and there was no significant difference between $FAP_{ORIGIN}$ or $FAP_{OPTIM}$ constructs in the percentage of non-fluorescent cells identified in these assays. Student's t-tests were used to compare $FAP_{OPTIM}$ to $FAP_{ORIGIN}$ for each construct (not significant = ns).

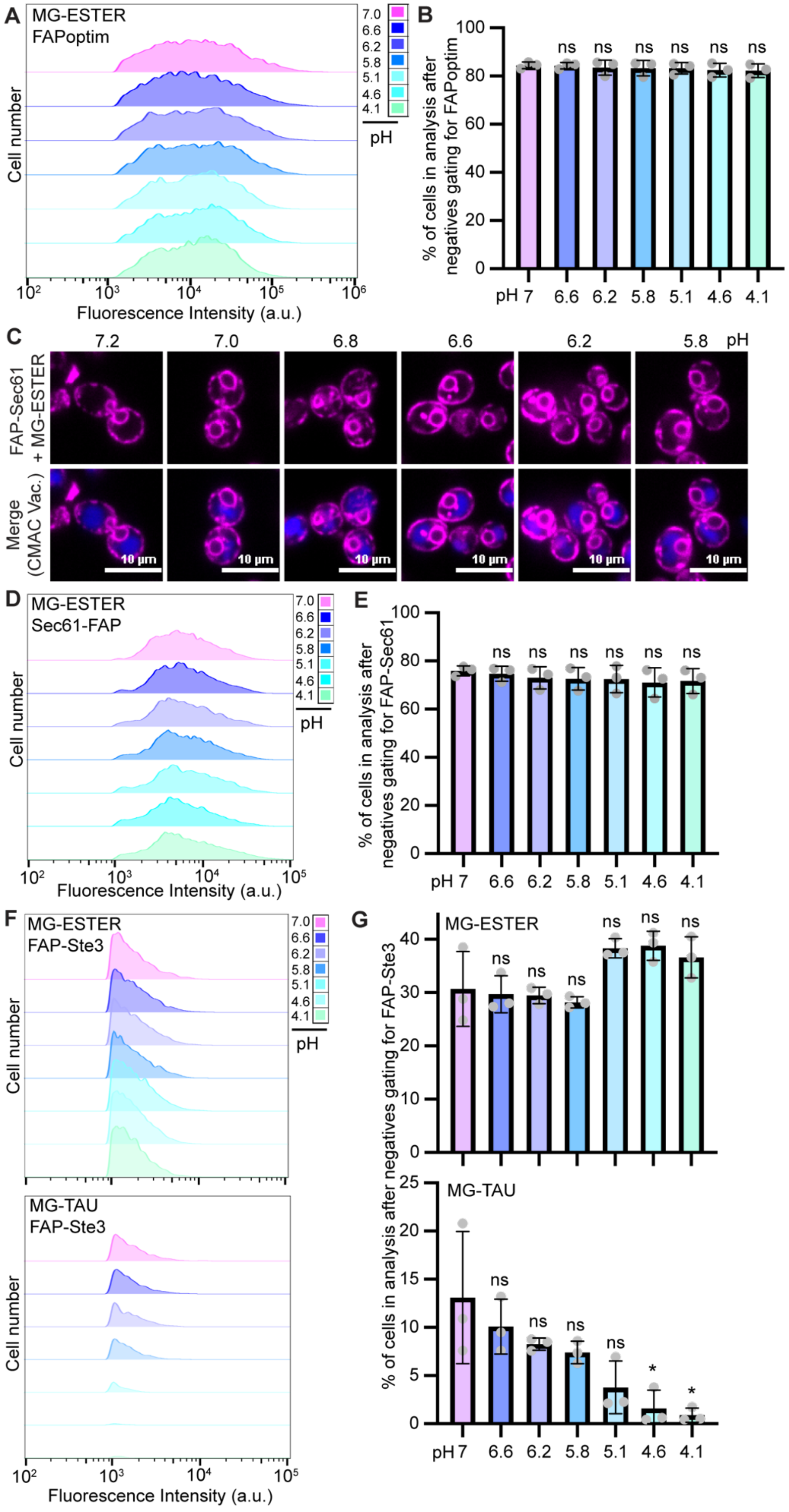

**Supplemental Figure S2 (accompanies Figure 3): FAP fluorescence response to pH changes.** (A and D) Histograms of cell cytometry data showing fluorescence distributions for cells exposed to the pHs indicated and expressing (A) FAP$_{\mathrm{OPTIM}}$ or (D) Sec61-FAP$_{\mathrm{OPTIM}}$. (B and E) The percentage of cells remaining in each experiment performed for Figure 2A-B (in panel B) or Figure 3C (in panel E) after the non-fluorescent cells were removed by gating. In all cases, ~80% of cells remained and there was no significant difference between FAP$_{\mathrm{OPTIM}}$ (panel B) or Sec61-FAP$_{\mathrm{OPTIM}}$ (panel E) at any pH in the percentage of non-fluorescent cells identified in these assays. Student's t-tests were used to compare FAP$_{\mathrm{OPTIM}}$ cells detected at lower pHs to those detected at pH 7.0 (not significant = ns). (C) Representative imaging for the data quantified by flow cytometry in panel 3C of the main paper, with $FAP_{OPTIM}$-Ste3 in magenta and CMAC in blue. (F) Histograms of flow cytometry data showing fluorescence distributions for cells exposed to the pHs indicated and expressing $FAP_{OPTIM}$-Ste3 and stained with either MG-ESTER (top) or MG-TAU (bottom). (G) The percentage of cells remaining in each experiment performed for Figure 3D-E after the non-fluorescent cells were removed by gating. While no significant pH-dependent changes were observed for MG-ESTER-$FAP_{OPTIM}$-Ste3 in the population of fluorescent cells (top), there was a dramatic loss in fluorescence for the MG-TAU incubated cells (bottom). Student's t-tests were used to compare $FAP_{OPTIM}$-Ste3 cells imaged at lower pH to those imaged at pH 7.0 (not significant = ns; $p < 0.05$ = *).

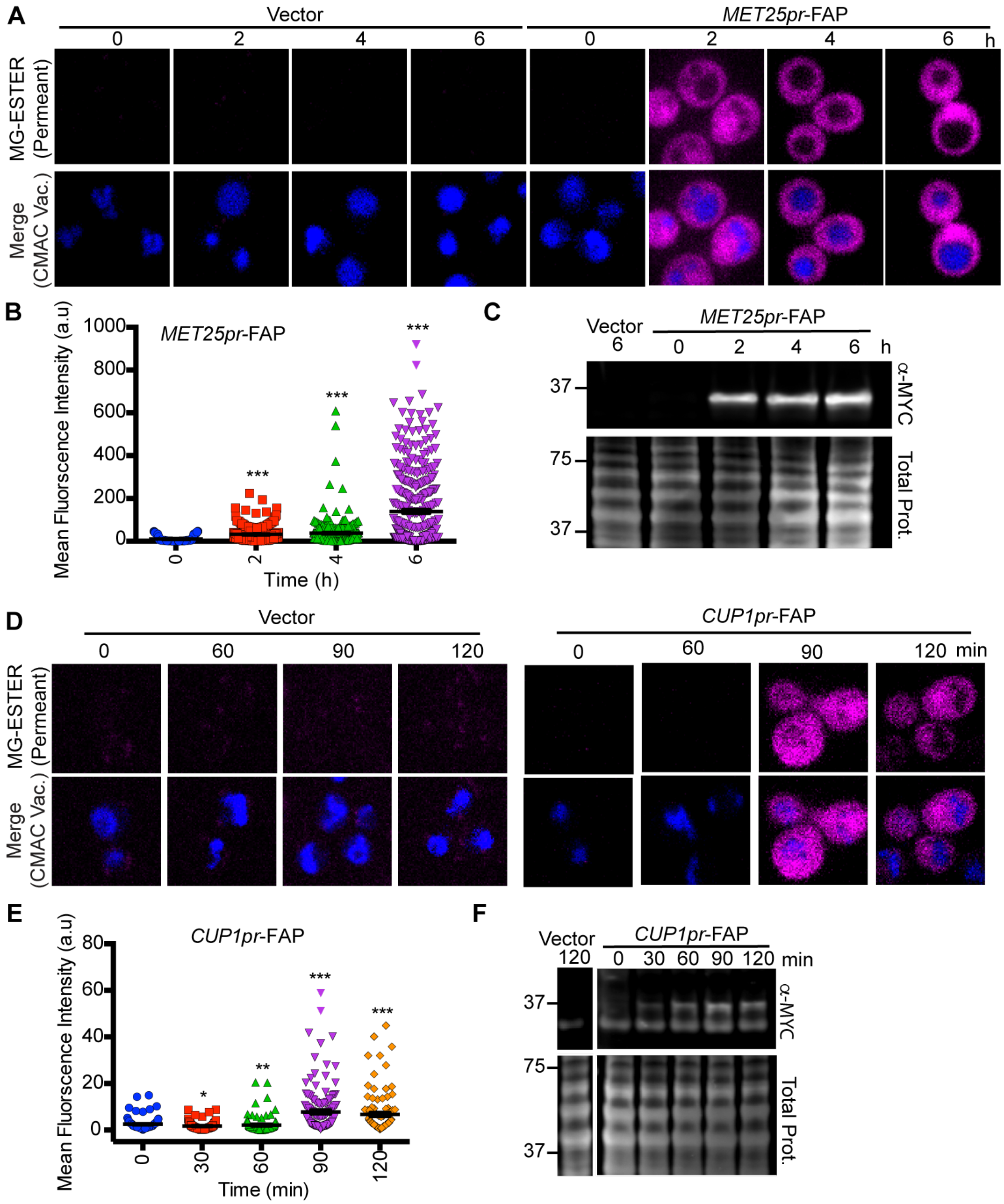

**Supplemental Figure S3 (accompanies Figure 4): Inducing FAP expression from *MET25* or *CUP1* promoters.** (A & D) Confocal fluorescence microscopy of soluble $FAP_{OPTIM}$ (magenta) expressed in WT cells from plasmids containing the *Met25pr* (A) or *Cup1pr* (D), with CMAC (blue) marking vacuoles. Images were captured at the times indicated and the empty vector control shows the background fluorescence when no FAP is expressed from these promoters. (B & E) Mean whole cell fluorescence intensity of $FAP_{OPTIM}$ signal from confocal microscopy in panels A and D, respectively. Kruskal-Wallis with Dunn's post hoc tests was performed, and statistical comparisons were made relative to the t=0 control. ($p<0.05$ = *; $p<0.005$ = **; $p<0.0005$ = ***). (C & F) Immunoblot of whole cell extracts from WT cells expressing the $FAP_{OPTIM}$ from the indicated promoter over time. MW markers are indicated on the left in kilodaltons.

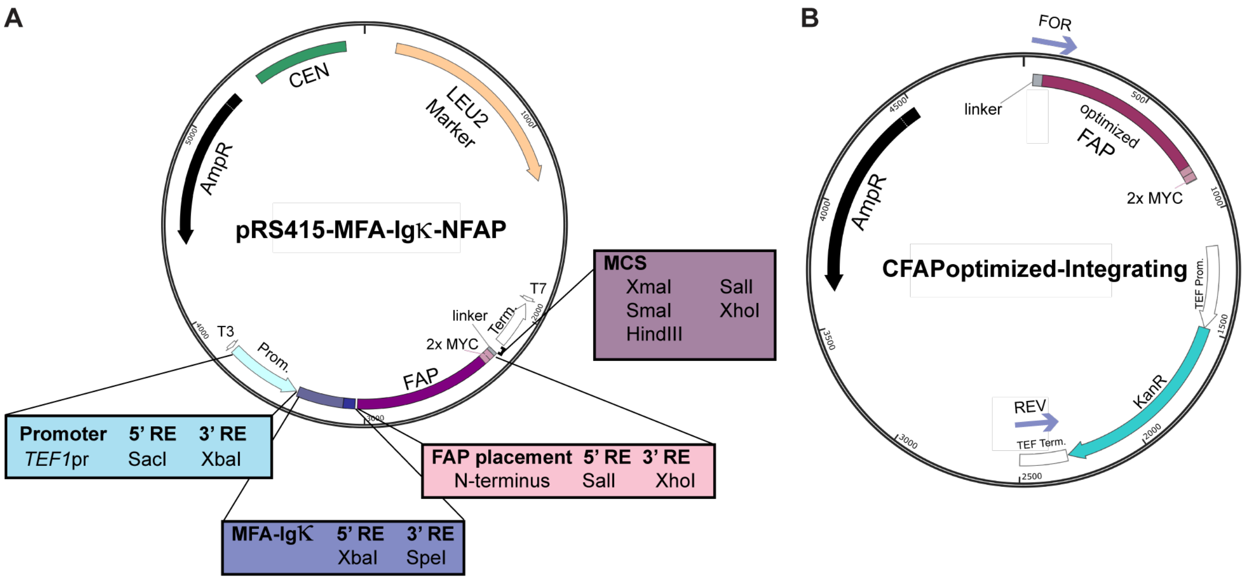

**Supplemental Figure S4 (accompanies Figure 4): FAP-expression and -tagging plasmids.** (A) Map of N-terminal FAP tagging plasmid with ER-targeting sequence from Mfa1 added to allow ER insertion. (B) Map of C-terminal FAP integration cassette marked with KANMX4. To amplify the FAP-KANMX4 cassette for C-terminal integration cloning use the F2 (indicated as FOR on map) and R1 (indicated as REV on map) primers, as described in (Longtine *et al.*, 1998), with gene-specific primer extensions added at the 5' end of each primer.

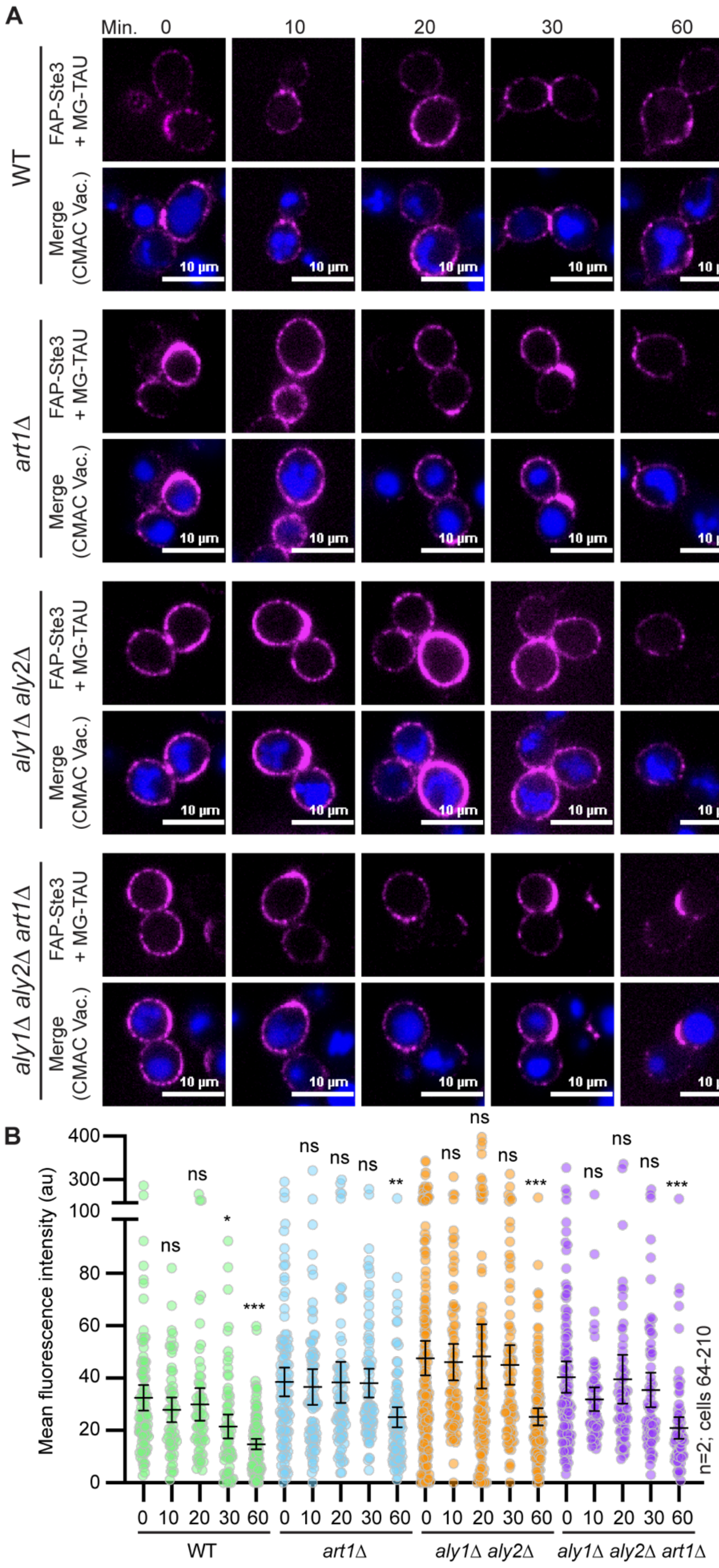

**Supplemental Figure S5 (accompanies Figure 9): Contribution of α-arrestins to steady-state turnover of FAP-Ste3.** (A) Representative confocal fluorescence microscopy images of $FAP_{OPTIM}$-Ste3 expressed from the endogenous *STE3pr* over time. Cells were incubated with MG-TAU (impermeant) dye at t=0 to visualize cell surface Ste3 (magenta) and CMAC (blue) stained the vacuoles. The dye was then washed from the cells before imaging at the indicated times, allowing us to monitor the steady-state turnover of FAP-Ste3 from the PM. (B) Kruskal-Wallis statistical analysis with Dunn's post hoc test was performed to compare the means of the three replicates to t=0 control for each of the strains (not significant = ns; $p < 0.05$ = *; $p < 0.005$ = **;$p < 0.0005$ = ***).

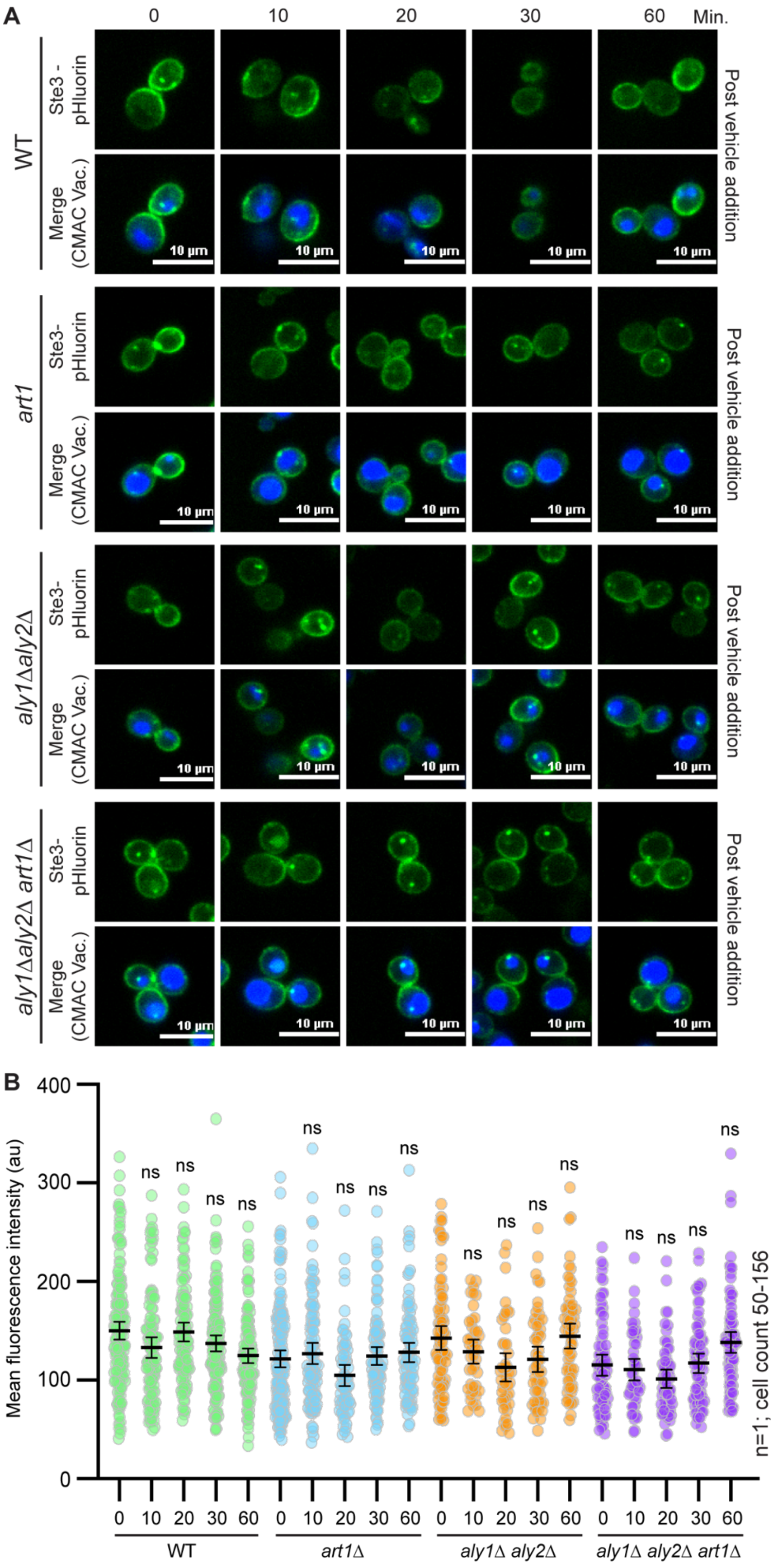

**Supplemental Figure S6 (accompanies Figure 10): Contribution of α-arrestins to steady-state turnover of Ste3-pHluorin.** (A-B) Representative confocal fluorescence microscopy images Ste3-pHluorin expressed from the *STE3* chromosomal locus (green), with CMAC (blue) staining to visualize vacuoles. Cells were imaged over time, without the addition of any ligand, and the mean whole cell fluorescence for the population was determined (B). Kruskal-Wallis statistical analysis with Dunn's post hoc test was performed to compare the means of the three replicates to t=0 control for each strain (not significant = ns).

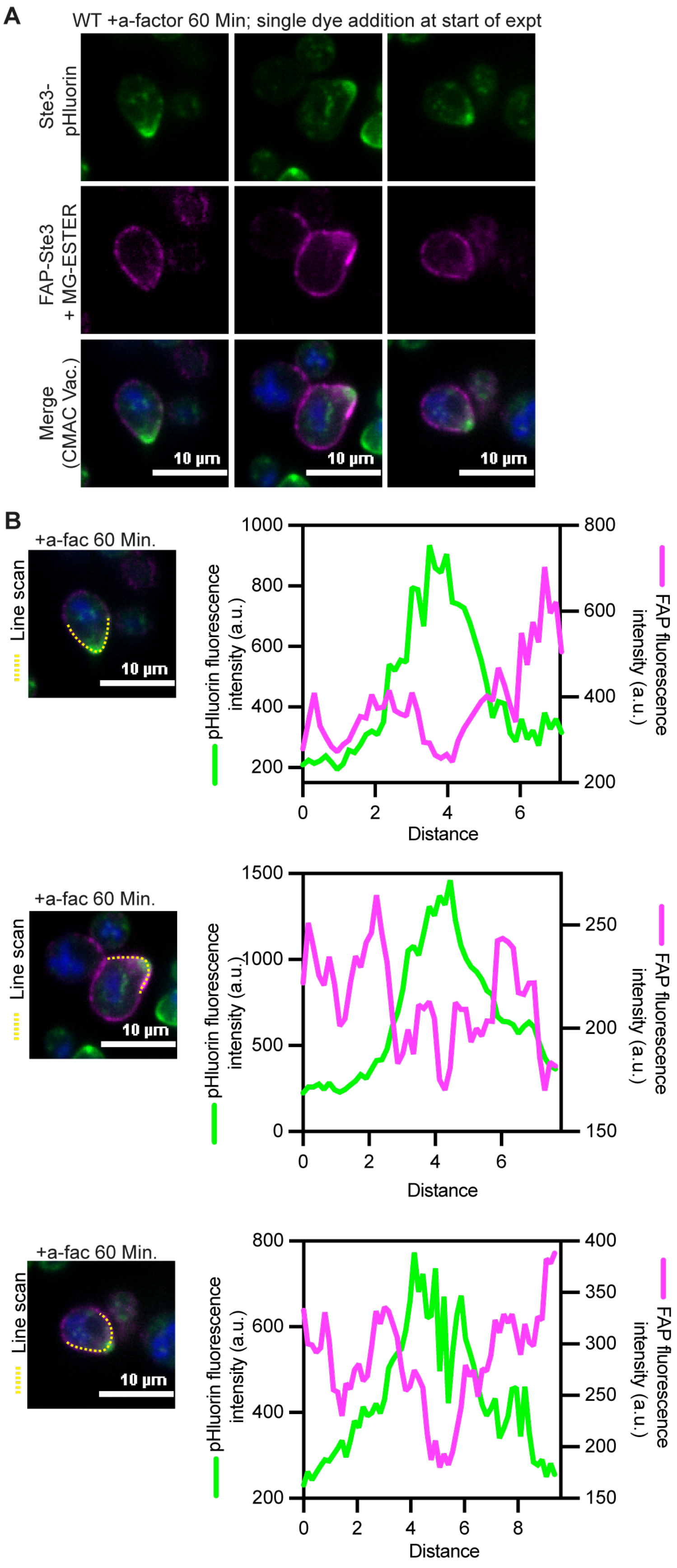

**Supplemental Figure S7 (accompanies Figure 11): FAP-Ste3 can be used to monitor recycling from the PM.** (A) Cells expressing $FAP_{OPTIM}$-Ste3 from a plasmid using the *STE3pr* and chromosomally integrated Ste3-pHluorin (green) were imaged by confocal fluorescence microscopy. Cells were incubated with MG-ESTER (magenta) and CMAC dye (blue). Once the MG-ESTER dye was washed from the cells, 5 $\mu$M of a-factor was added, and cells were imaged at 60 min with no further dye additions. This allowed us to monitor a single pool of Ste3 at the onset of the experiment with FAP while monitoring the total pool of Ste3 with pHluorin. These images are added in support of the data shown in Figure 11C. (B) Line scan analysis to monitor the relative distributions of FAP-Ste3 and Ste3-pHluorin after pheromone addition. The region used in the line scan is indicated as a dashed yellow line on the image and the fluorescence intensities along this line for GFP (left y-axis) or FAP (right y-axis) are plotted. In each case, the Ste3-pHluorin is distributed to the bud tip while the $FAP_{OPTIM}$-Ste3 localizes to the regions adjacent to the shmoo tip.

**Supplemental Table 1. Yeast strains used in this study.**

| Strain | Genotype | Reference or Description |
|---|---|---|
| BY4742 | *MAT* a *his3*Δ*1 leu2*Δ*0 lys2*Δ*0 ura3*Δ*0* | (Brachmann *et al.*, 1998) |
| *yps1*Δ | *MAT* a *his3*Δ*1 leu2*Δ*0 lys2*Δ*0 ura3*Δ*0 ysp1*Δ::KAN | (Winzeler *et al.*, 1999) |
| *mkc7*Δ | *MAT* a *his3*Δ*1 leu2*Δ*0 lys2*Δ*0 ura3*Δ*0 mkc7*Δ*::HPHMX4* | The *MCK7* coding sequence was replaced with the hygromycin resistance cassette using the (Longtine *et al.*, 1998) strategy in the BY4742 background. |
| *yps1*Δ *mkc7*Δ | *MAT* a *his3*Δ*1 leu2*Δ*0 lys2*Δ*0 ura3*Δ*0 ysp1*Δ::KAN *mkc7*Δ*::HPHMX4* | The *MCK7* coding sequence was replaced with the hygromycin resistance cassette using the (Longtine *et al.*, 1998) strategy in the *yps1*Δ::KAN background. |
| *aly1*Δ *aly2*Δ *mkc7*Δ | *MAT* a *his3*Δ*1 leu2*Δ*0 lys2*Δ*0 ura3*Δ*0 aly1*Δ*::KAN aly2*Δ*::KAN mkc7*Δ*::HPHMX4* | The *MCK7* coding sequence was replaced with the hygromycin resistance cassette using the (Longtine *et al.*, 1998) strategy in the *aly1*Δ::KAN *aly2*Δ::KAN background from (O'Donnell *et al.*, 2010) |
| *art1*Δ *mkc7*Δ | *MAT* a *his3*Δ*1 leu2*Δ*0 lys2*Δ*0 ura3*Δ*0 art1*Δ*::NAT mkc7*Δ*::HPHMX4* | This study. The *MCK7* coding sequence was replaced with the hygromycin resistance cassette using the (Longtine *et al.*, 1998) strategy in the *art1*Δ::KAN background. |
| *aly1*Δ *aly2*Δ *art1*Δ *mkc7*Δ | *MAT* a *his3*Δ*1 leu2*Δ*0 lys2*Δ*0 ura3*Δ*0 aly1*Δ*::NAT aly2*Δ*::URA3 art1*Δ*::KAN mkc7*Δ*::HPHMX4* | The *MCK7* coding sequence was replaced with the hygromycin resistance cassette using the (Longtine *et al.*, 1998) strategy in the *aly1*Δ::KAN *aly2*Δ::KAN background from Prosser et al. |
| *ste3*Δ | *MAT* a *his3*Δ*1 leu2*Δ*0 lys2*Δ*0 ura3*Δ*0* ste3Δ::KAN | (Winzeler et al., 1999) |
| Ste3-pHluorin | *MAT* a *his3*Δ*1 leu2*Δ*0 lys2*Δ*0 ura3*Δ*0 STE3-pHlourin::HPHMX4* | Ste3 was tagged at its chromosomal locus using a PCR-based strategy where the pHluorin::*HPHMX4* cassette from the pDP0437 plasmid (see below and Figure S4) was amplified and this was transformed into the BY4742 background. Correct location of integrations was validated by PCR. |

| Strain | Genotype | Reference or Description |
|---|---|---|
| *art1Δ* Ste3-pHluorin | *MAT* a *his3Δ1 leu2Δ0 lys2Δ0 ura3Δ0 art1Δ::NAT Ste3-pHluorin:HPHMX4* | Ste3 was tagged at its chromosomal locus using a PCR-based strategy where the pHluorin::*HPHMX4* cassette from the pDP0437 plasmid (see below and Figure S4) was amplified and this was transformed into the *art1Δ* background (Winzeler *et al.*, 1999). Correct location of integrations was validated by PCR. |
| *aly1Δ aly2Δ* Ste3-pHluorin | *MAT* a *his3Δ1 leu2Δ0 lys2Δ0 ura3Δ0 aly1Δ::KAN aly2Δ::KAN Ste3-pHluorin::HPHMX4* | Ste3 was tagged at its chromosomal locus using a PCR-based strategy where the pHluorin::*HPHMX4* cassette from the pDP0437 plasmid (see below and Figure S4) was amplified and this was transformed into the *aly1Δ aly2Δ*background (O'Donnell *et al.*, 2010). Correct location of integrations was validated by PCR. |
| *aly1Δ aly2Δ art1Δ* Ste3-pHluorin | *MAT* a *his3Δ1 leu2Δ0 lys2Δ0 ura3Δ0 aly1Δ::KAN aly2Δ::KAN art1Δ::NAT Ste3-pHluorin::HPHMX4* | Ste3 was tagged at its chromosomal locus using a PCR-based strategy where the pHluorin::*HPHMX4* cassette from the pDP0437 plasmid (see below and Figure S4) was amplified and this was transformed into the *aly1Δ aly2Δ art1Δ* background (Prosser *et al.*, 2015). Correct location of integrations was validated by PCR |
| 9ArrΔ Ste3-pHluorin | *MAT a his3Δ1 leu2Δ0 lys2Δ0 ura3Δ0 aly1Δ0 aly2Δ0 art1Δ0 rod1Δ::NAT rog3Δ::NAT ecm21Δ::KAN csr2Δ::KAN art5Δ::HIS3 Ste3-pHlourin::HPHMX* | EN60 strain (a.k.a. 9ArrΔ) made in (Nikko and Pelham, 2009), was modified in this study to include the Ste3-pHluorin integration. |

**Supplemental Table 2. Plasmids used in this study.**

| Plasmid name | Genotype | Reference or Description | Addgene ID |
|---|---|---|---|
| pRS415-TEF1pr | *TEF1pr* CEN *LEU2* | (Mumberg *et al.*, 1995) | |
| pRS415-TEF1pr-$NFAP_{orig}$ | *TEF1pr*- $NFAP_{orig}$ CEN *LEU2* | The FAPorig-MYC-Linker sequence was cloned into the *Bam*HI and *Sma*I sites of the pRS415-TEF1pr plasmid. | 221086 |
| pRS415-TEF1pr-$CFAP_{orig}$ | *TEF1pr*- $CFAP_{orig}$ CEN *LEU2* | The Linker-MYC-FAPorig-MYC sequence was cloned into the *Sal*I and *Xho*I sites of the pRS415-TEF1pr plasmid. | 221087 |
| pRS415-TEF1pr-$NFAP_{optim}$ | *TEF1pr*- $NFAP_{optim}$ CEN *LEU2* | The FAPoptim-MYC-Linker sequence was cloned into the *Bam*HI and *Sma*I sites of the pRS415-TEF1pr plasmid. | 221088 |
| pRS415-TEF1pr-$CFAP_{optim}$ | *TEF1pr*- $CFAP_{optim}$ CEN *LEU2* | The Linker-MYC-FAPoptim-MYC sequence was cloned into the *Sal*I and *Xho*I sites of the pRS415-TEF1pr plasmid. | 221089 |
| pRS415-TEF1pr-MFA-$NFAP_{optim}$ | *TEF1pr*-MFA1 signal sequence-$NFAP_{optim}$ CEN *LEU2* | The FAPoptim-MYC-Linker sequence was cloned into the *Bam*HI and *Sma*I sites of the pRS415-TEF1pr plasmid and the *MFA1* signal sequence was cloned into the *Xba*I/*Spe*I sites upstream of the FAP tag to help target constructs to the ER. | 221090 |
| pRS413-TEF1pr | *TEF1pr* CEN *HIS3* | (Mumberg *et al.*, 1995) | |
| pRS413-TEF1pr-$NFAP_{orig}$ | *TEF1pr*- $NFAP_{orig}$ CEN *HIS3* | The FAPorig-MYC-Linker sequence was cloned into the *Bam*HI and *Sma*I sites of the pRS413-TEF1pr plasmid. | 221091 |
| pRS413-TEF1pr-$CFAP_{orig}$ | *TEF1pr*- $CFAP_{orig}$ CEN *HIS3* | The Linker-MYC-FAPorig-MYC sequence was cloned into the *Sal*I and *Xho*I sites of the pRS413-TEF1pr plasmid. | 221092 |
| pRS413-TEF1pr-$NFAP_{optim}$ | *TEF1pr*- $NFAP_{optim}$ CEN *HIS3* | The FAPoptim-MYC-Linker sequence was cloned into the *Bam*HI and *Sma*I sites of the pRS413-TEF1pr plasmid. | 221093 |
| pRS413-TEF1pr-$CFAP_{optim}$ | *TEF1pr*- $CFAP_{optim}$ CEN *HIS3* | The Linker-MYC-FAPoptim-MYC sequence was cloned into the *Sal*I and *Xho*I sites of the pRS413-TEF1pr plasmid. | 221094 |
| pRS413-TEF1pr-MFA-$NFAP_{optim}$ | *TEF1pr*-MFA1 signal sequence-$NFAP_{optim}$ CEN *HIS3* | The FAPoptim-MYC-Linker sequence was cloned into the *Bam*HI and *Sma*I sites of the pRS413-TEF1pr plasmid and the *MFA1* signal sequence was cloned into the *Xba*I/*Spe*I sites upstream of the FAP tag to help target constructs to the ER. | 221095 |

| Plasmid name | Genotype | Reference or Description | Addgene ID |
|---|---|---|---|
| pRS415-ADH1pr | *ADH1pr* CEN *LEU2* | (Mumberg *et al.*, 1995) | |
| pRS415-ADH1pr-$NFAP_{optim}$ | *ADH1pr*- $NFAP_{optim}$ CEN *LEU2* | The FAPoptim-MYC-Linker sequence was cloned into the *Bam*HI and *Sma*I sites of the pRS415-ADH1pr plasmid. | 221096 |
| pRS415-ADH1pr-$CFAP_{optim}$ | *ADH1p*- $CFAP_{optim}$ CEN *LEU2* | The Linker-MYC-FAPoptim-MYC sequence was cloned into the *Sal*I and *Xho*I sites of the pRS415-ADH1pr plasmid. | 221097 |
| pRS413-ADH1pr | *ADH1pr* CEN *HIS3* | (Mumberg *et al.*, 1995) | |
| pRS413-ADH1pr-$NFAP_{optim}$ | *ADH1pr*- $NFAP_{optim}$ CEN *HIS3* | The FAPoptim-MYC-Linker sequence was cloned into the *Bam*HI and *Sma*I sites of the pRS413-ADH1pr plasmid. | 221098 |
| pRS413-ADH1pr-$CFAP_{optim}$ | *ADH1pr*- $CFAP_{optim}$ CEN *HIS3* | The Linker-MYC-FAPoptim-MYC sequence was cloned into the *Sal*I and *Xho*I sites of the pRS413-ADH1pr plasmid. | 221099 |
| pRS415-MET25pr | *MET25pr* CEN *LEU2* | The *TEF1* promoter from the pRS415-TEF1pr plasmid (Mumberg *et al.*, 1995) was removed by restriction digestion and replaced with the *MET25* promoter (PCR amplified from the pUG vector series) flanked by *Sac*I and *Xba*I restriction sites. | 221100 |
| pRS415-MET25pr-$NFAP_{optim}$ | *MET25pr* CEN *LEU2* | The FAPoptim-MYC-Linker sequence was cloned into the *Bam*HI and *Sma*I sites of the pRS415-MET25pr plasmid. | 221101 |
| pRS415-MET25pr-$CFAP_{optim}$ | *MET25pr* CEN *LEU2* | The Linker-MYC-FAPoptim-MYC sequence was cloned into the *Sal*I and *Xho*I sites of the pRS415-MET25pr plasmid. | 221102 |
| pRS413-MET25pr | *MET25pr* CEN *HIS3* | The *TEF1* promoter from the pRS413-TEF1pr plasmid (Mumberg *et al.*, 1995) was removed by restriction digestion and replaced with the *MET25* promoter (PCR amplified from the pUG vector series) flanked by *Sac*I and *Xba*I restriction sites. | 221103 |
| pRS413-MET25pr-$NFAP_{optim}$ | *MET25pr* CEN *HIS3* | The $FAP_{optim}$-MYC-Linker sequence was cloned into the *Bam*HI and *Sma*I sites of the pRS413-MET25pr plasmid. | 221104 |

| Plasmid name | Genotype | Reference or Description | Addgene ID |
|---|---|---|---|
| pRS413-MET25pr-CFAP$_{optim}$ | *MET25pr* CEN *HIS3* | The Linker-MYC-FAP$_{optim}$-MYC sequence was cloned into the *Sal*I and *Xho*I sites of the pRS413-MET25pr plasmid. | 221105 |
| pRS415-CUP1pr | *CUP1pr* CEN *LEU2* | The *TEF1* promoter from the pRS415-TEF1pr plasmid (Mumberg *et al.*, 1995) was removed by restriction digestion and replaced with the *CUP1* promoter (PCR amplified from the pKK212 plasmid used in (O'Donnell *et al.*, 2013) flanked by *Sac*I and *BamH*I restriction sites. | 221106 |
| pRS415-CUP1pr-NFAP$_{optim}$ | *CUP1pr*- NFAP$_{optim}$ CEN *LEU2* | The FAPoptim-MYC-Linker sequence was cloned into the *Bam*HI and *Sma*I sites of the pRS415-CUP1pr plasmid. | 221107 |
| pRS415-CUP1pr-CFAP$_{optim}$ | *CUP1pr*- CFAP$_{optim}$ CEN *LEU2* | The Linker-MYC-FAP$_{optim}$-MYC sequence was cloned into the *Sal*I and *Xho*I sites of the pRS415-CUP1pr plasmid. | 221108 |
| pRS413-CUP1pr | *CUP1pr* CEN *HIS3* | The *TEF1* promoter from the pRS413-TEF1pr plasmid (Mumberg *et al.*, 1995) was removed by restriction digestion and replaced with the *CUP1* promoter (PCR amplified from the pKK212 plasmid used in (O'Donnell *et al.*, 2013) flanked by *Sac*I and *BamH*I restriction sites. | 221109 |
| pRS413-CUP1pr-NFAP$_{optim}$ | *CUP1pr*- NFAP$_{optim}$ CEN *HIS3* | The FAP$_{optim}$-MYC-Linker sequence was cloned into the *Bam*HI and *Sma*I sites of the pRS413-CUP1pr plasmid. | 221110 |
| pRS413-CUP1pr-CFAP$_{optim}$ | *CUP1pr*- CFAP$_{optim}$ CEN *HIS3* | The Linker-MYC-FAP$_{optim}$-MYC sequence was cloned into the *Sal*I and *Xho*I sites of the pRS413-CUP1pr plasmid. | 221111 |
| pRS415-*STE3pr-MFA-Igkappa-FAP-STE3* | *TEF1pr* CEN *LEU2* | The Ste3 promoter sequence (500 bprs upstream of the Ste3 ATG) was amplified from chromosomal DNA extracts using primers containing restriction enzyme adaptors for *Xho*I and *Xba*I. The pRS415-*TEF1pr*-Ste3-GFP plasmid was then digested with *Xho*I/*Xba*I to remove the *TEF1* promoter and the Ste3 promoter sequences was ligated into its place. | 221112 |
| pRS415 | CEN *LEU2* | (Sikorski and Hieter, 1989) | |

| Plasmid name | Genotype | Reference or Description | Addgene ID |
|---|---|---|---|
| pRS415-*TEF1pr-STE3-GFP* | *Ste3pr* CEN *LEU2* | The pRS415-*TEF1pr* plasmid (Mumberg *et al.*, 1995) was used to insert the Ste3 coding sequence fused at the C-terminus to GFP. The GFP was inserted into the plasmid first between the *Xho*I and *Sal*I restriction enzyme sites and then Ste3 coding sequence (lacking the stop codon) was inserted into the *Spe*I and *Sma*I restriction enzyme sites. | 221113 |
| pRS415-*TEF1pr-MFA-Igkappa-FAP-STE3* | *TEF1pr* CEN LEU2 | The pRS415-TEF1pr-MFA-NFAP$_{optim}$ plasmid (described above) was used to insert the Ste3 coding sequence as an in-frame N-terminal fusion to FAP. The Ste3 coding sequence (lacking the stop codon) was inserted into the *Sma*I and *Xho*I restriction enzyme sites. The resulting plasmid contains the *MFA1* ER targeting and secretion sequence, an N-terminal FAP$_{optim}$ tag on Ste3. | 221114 |
| pRS415-*TEF1pr-FAP$_{orig}$-SEC61* | *TEF1pr* CEN LEU2 | (Hager *et al.*, 2018) | |
| pRS415-*TEF1pr-FAP$_{orig}$-SEC63* | *TEF1pr* CEN LEU2 | (Hager *et al.*, 2018) | |
| pRS415-*TEF1pr-FAP$_{optim}$-SEC61* | *TEF1pr* CEN LEU2 | The FAP$_{origin}$ sequence in the SEC61 plasmid was replaced with FAP$_{optim}$ using restriction digestion. | 221115 |
| pRS415-*TEF1pr-FAP$_{optim}$-SEC63* | *TEF1pr* CEN LEU2 | The FAP$_{origin}$ sequence in the SEC61 plasmid was replaced with FAP$_{optim}$ using restriction digestion. | 221116 |
| pRS415-*STE3pr-MFA-Igkappa-FAP-STE3-TAILLESS* | *STE3pr-FAP-STE3-Tailless* CEN LEU2 | PCR amplification of the STE3 coding sequence lacking amino acids 288-end (the C-tail of Ste3), where amino acid 288 was converted to a stop codon via PCR, was preformed and this DNA was subcloned into pRS415-*STE3pr-MFA-Igkappa-FAP* plasmid at the *Sma*I and *Xho*I sites. | 221117 |
| pRS415-*STE3pr-MFA-Igkappa-FAP-STE3-ΔPEST* | *STE3pr-FAP-STE3-ΔPEST* CEN LEU2 | Amino acid 413 was mutated to a stop codon using site-directed mutagenesis, creating a truncation mutant of Ste3's C-tail at this residue in the pRS415-*STE3pr-MFA-Igkappa-FAP-STE3.* | 221118 |
| pRS415-*STE3pr-MFA-Igkappa-FAP-STE3-K424R* | *STE3pr-FAP-STE3-K424R* CEN LEU2 | The lysine at position 424 in Ste3's C-tail was mutated to arginine using site-directed mutagenesis of pRS415-*STE3pr-MFA-Igkappa-FAP-STE3.* | 221110 |

| Plasmid name | Genotype | Reference or Description | Addgene ID |
|---|---|---|---|
| pRS413-TEF1pr-ANP1-NFAP | *TEF1pr-ANP1-NFAP* CEN HIS3 | The coding sequence of *ANP1* lacking the stop codon was PCR amplified with primers containing *Bam*HI and *Sma*I restriction enzyme adaptors. The PCR product was then inserted into the pRS413-TEF1pr-$NFAP_{optim}$ plasmid using standard cloning approaches. | 221120 |
| pRS413-TEF1pr-ERG6-NFAP | *TEF1pr-* ERG6-NFAP CEN HIS3 | The coding sequence of *ERG6* lacking the stop codon was PCR amplified with primers containing *Bam*HI and *Sma*I restriction enzyme adaptors. The PCR product was then inserted into the pRS413-TEF1pr-$NFAP_{optim}$ plasmid using standard cloning approaches. | 221121 |
| pRS413-TEF1pr-PMA1-NFAP | *TEF1pr-* PMA1-NFAP CEN HIS3 | The coding sequence of *PMA1* lacking the stop codon was PCR amplified with primers containing *Bam*HI and *Sma*I restriction enzyme adaptors. The PCR product was then inserted into the pRS413-TEF1pr-$NFAP_{optim}$ plasmid using standard cloning approaches. | 221122 |
| pRS413-TEF1pr-RPA34-NFAP | *TEF1pr-* RPA34-NFAP CEN HIS3 | The coding sequence of *RPA34* lacking the stop codon was PCR amplified with primers containing *Bam*HI and *Sma*I restriction enzyme adaptors. The PCR product was then inserted into the pRS413-TEF1pr-$NFAP_{optim}$ plasmid using standard cloning approaches. | 221123 |
| pRS413-TEF1pr-SEC7-NFAP | *TEF1pr-* SEC7-NFAP CEN HIS3 | The coding sequence of *SEC7* lacking the stop codon was PCR amplified with primers containing *Bam*HI and *Sma*I restriction enzyme adaptors. The PCR product was then inserted into the pRS413-TEF1pr-$NFAP_{optim}$ plasmid using standard cloning approaches. | 221124 |
| pRS413-TEF1pr-VPH1-NFAP | *TEF1pr-* VPH1-NFAP CEN HIS3 | The coding sequence of *VPH1* lacking the stop codon was PCR amplified with primers containing *Bam*HI and *Sma*I restriction enzyme adaptors. The PCR product was then inserted into the pRS413-TEF1pr-$NFAP_{optim}$ plasmid using standard cloning approaches. | 221125 |

| Plasmid name | Genotype | Reference or Description | Addgene ID |
|---|---|---|---|
| pRS413-TEF1pr-SEC61-NFAP | *TEF1pr-* SEC61-NFAP CEN HIS3 | The coding sequence of *SEC61* lacking the stop codon was PCR amplified with primers containing *Bam*HI and *Sma*I restriction enzyme adaptors. The PCR product was then inserted into the pRS413-TEF1pr-$NFAP_{optim}$ plasmid using standard cloning approaches. | 221126 |
| pDP0437 | pFA6-pHluorin::*HPHMX* | This pHluorin integration plasmid was constructed by subcloning the 977 bp SalI/BglII fragment from pBW1571 (pFA6a-pHluorin::*KANMX6* from (Prosser *et al.*, 2010) into the SalI/BglII sites of pAG32 (pFA6-HPHMX4 described in (Goldstein and McCusker, 1999). The resulting plasmid was used for C-terminal tagging of the endogenous *STE3* locus using a PCR-based integration strategy described in (Longtine *et al.*, 1998) | 221127 |
| pFA6-FAPc-KAN | pFA6-FAPoptim::*KANMX* | This FAPoptim integration plasmid was constructed by subcloning the FAPoptim sequence into the pFA6a-KANMX4 plasmid described in (Goldstein and McCusker, 1999). The resulting plasmid can be used for C-terminal tagging of endogenous protiens using the PCR-based integration strategy described in (Longtine *et al.*, 1998). | 221128 |
| pFA6-FAPc-NAT | pFA6-FAPoptim::*NATMX* | This FAPoptim integration plasmid was constructed by subcloning the FAPoptim sequence into the pFA6a-NATMX4 plasmid described in (Goldstein and McCusker, 1999). The resulting plasmid can be used for C-terminal tagging of endogenous protiens using the PCR-based integration strategy described in (Longtine *et al.*, 1998). | 221129 |

**Supplemental References**

Brachmann, C.B., Davies, A., Cost, G.J., Caputo, E., Li, J., Hieter, P., and Boeke, J.D. (1998). Designer deletion strains derived from Saccharomyces cerevisiae S288C: a useful set of strains and plasmids for PCR-mediated gene disruption and other applications. Yeast *14*, 115-132.

Goldstein, A.L., and McCusker, J.H. (1999). Three new dominant drug resistance cassettes for gene disruption in Saccharomyces cerevisiae. Yeast *15*, 1541-1553.

Hager, N.A., Krasowski, C.J., Mackie, T.D., Kolb, A.R., Needham, P.G., Augustine, A.A., Dempsey, A., Szent-Gyorgyi, C., Bruchez, M.P., Bain, D.J., Kwiatkowski, A.V., O'Donnell, A.F., and Brodsky, J.L. (2018). Select alpha-arrestins control cell-surface abundance of the mammalian Kir2.1 potassium channel in a yeast model. J Biol Chem *293*, 11006-11021.

Longtine, M.S., McKenzie, A., 3rd, Demarini, D.J., Shah, N.G., Wach, A., Brachat, A., Philippsen, P., and Pringle, J.R. (1998). Additional modules for versatile and economical PCR-based gene deletion and modification in Saccharomyces cerevisiae. Yeast *14*, 953-961.

Mumberg, D., Muller, R., and Funk, M. (1995). Yeast vectors for the controlled expression of heterologous proteins in different genetic backgrounds. Gene *156*, 119-122.

Nikko, E., and Pelham, H.R. (2009). Arrestin-mediated endocytosis of yeast plasma membrane transporters. Traffic *10*, 1856-1867.

O'Donnell, A.F., Apffel, A., Gardner, R.G., and Cyert, M.S. (2010). Alpha-arrestins Aly1 and Aly2 regulate intracellular trafficking in response to nutrient signaling. Mol Biol Cell *21*, 3552-3566.

O'Donnell, A.F., Huang, L., Thorner, J., and Cyert, M.S. (2013). A calcineurin-dependent switch controls the trafficking function of alpha-arrestin Aly1/Art6. J Biol Chem *288*, 24063-24080.

Prosser, D.C., Pannunzio, A.E., Brodsky, J.L., Thorner, J., Wendland, B., and O'Donnell, A.F. (2015). alpha-Arrestins participate in cargo selection for both clathrin-independent and clathrin-mediated endocytosis. J Cell Sci *128*, 4220-4234.

Prosser, D.C., Whitworth, K., and Wendland, B. (2010). Quantitative analysis of endocytosis with cytoplasmic pHluorin chimeras. Traffic *11*, 1141-1150.

Sikorski, R.S., and Hieter, P. (1989). A system of shuttle vectors and yeast host strains designed for efficient manipulation of DNA in Saccharomyces cerevisiae. Genetics *122*, 19-27.

Winzeler, E.A., Shoemaker, D.D., Astromoff, A., Liang, H., Anderson, K., Andre, B., Bangham, R., Benito, R., Boeke, J.D., Bussey, H., Chu, A.M., Connelly, C., Davis, K., Dietrich, F., Dow, S.W., El Bakkoury, M., Foury, F., Friend, S.H., Gentalen, E., Giaever, G., Hegemann, J.H., Jones, T., Laub, M., Liao, H., Liebundguth, N., Lockhart, D.J., Lucau-Danila, A., Lussier, M., M'Rabet, N., Menard, P., Mittmann, M., Pai, C.,

Rebischung, C., Revuelta, J.L., Riles, L., Roberts, C.J., Ross-MacDonald, P., Scherens, B., Snyder, M., Sookhai-Mahadeo, S., Storms, R.K., Veronneau, S., Voet, M., Volckaert, G., Ward, T.R., Wysocki, R., Yen, G.S., Yu, K., Zimmermann, K., Philippsen, P., Johnston, M., and Davis, R.W. (1999). Functional characterization of the S. cerevisiae genome by gene deletion and parallel analysis. Science *285*, 901-906.
